# Assessing multi-eGO Predictions of PDZ2–Peptide Binding across Mutations

**DOI:** 10.64898/2026.09.10.750618

**Authors:** Camilla Ardizzone, Bruno Stegani, Fran Bačić Toplek, Stefano Gianni, Riccardo Capelli, Carlo Camilloni

**Author notes:** Corresponding authors: Riccardo Capelli, Carlo Camilloni. CNRS, Ecole Normale Supérieure de Lyon, Laboratoire de Physique, UMR5672, Lyon 69342, France.

## Abstract

Accurately predicting how mutations alter protein–peptide binding remains challenging for molecular simulations because both conformational sampling and binding kinetics are computationally demanding. Here, we investigate whether multi-eGO, a hybrid transferable/structure-based atomistic-resolution model previously developed and validated for protein–small molecule interactions, can be transferred to protein–peptide binding. Using the PDZ2 domain of protein tyrosine phosphatase basophil-like in complex with the peptide EQVTAV as a benchmark, we show that multi-eGO, calibrated on explicit-solvent simulations of the complex, predicts the equilibrium dissociation constant of the complex within the expected uncertainty, although binding and unbinding are both accelerated by several orders of magnitude. We then introduce conservative mutations in PDZ2 and in the peptide without additional training for each mutant. For the PDZ2 mutants in complex with EQVTAV, the predicted dissociation constants follow experiment (Pearson r = 0.97), with a compressed response, whereas smaller effects, such as that of the peptide mutation and the coupling between peptide and protein mutations, are not resolved. A maximum-caliber reweighting of the simulated first-passage times shows that a single acceleration factor, common to all twelve complexes, accounts for most of the kinetic discrepancy, and that the remaining system-specific errors in the rate constants are recovered by reweighting the simulated events. Up to a global time scale, the binding and unbinding events sampled by multi-eGO are thus consistent with the experimental kinetics. These results define multi-eGO as a computationally efficient approach to estimate the effects of conservative protein mutations on peptide binding affinity and to interpret the associated kinetics, while absolute rate constants remain out of reach.

## INTRODUCTION

Protein–peptide interactions mediate a large fraction of cellular signaling, and their quantitative characterization, including binding affinity and the kinetics of complex formation and dissociation, is central to understanding processes ranging from signal transduction to protein trafficking^1,2^. PDZ (PSD-95/Discs-large/ZO-1) domains are among the most common protein–protein interaction modules found in nature. They recognize short linear motifs, typically located at the C-terminus of their partner proteins, and are frequently found in tandem or within scaffolding proteins that organize multiprotein signaling complexes^3,4^. Because PDZ–peptide interactions are promiscuous and can be modulated by individual amino acid substitutions^5^, they provide an attractive system for testing how well computational models capture the subtle energetic determinants of protein–peptide recognition.

Molecular dynamics (MD) simulations offer, in principle, a route to dissecting the thermodynamics and kinetics of such interactions at atomic resolution. However, converged sampling of binding and unbinding events for protein–peptide complexes remains computationally demanding with all-atom, explicit-solvent models, because these events are rare on the timescales accessible even to well-resourced simulations^6,7^. One approach to overcoming this limitation is provided by enhanced sampling methods, which can accelerate the exploration of conformational space by introducing external biases or modified sampling protocols and can be used to characterize thermodynamics, kinetics, or, in some cases, both^8–11^. Their application, however, often requires careful identification of the relevant degrees of freedom and states associated with the process of interest. Simplified models, such as coarse-grained and structure-based models, provide an alternative by reducing the computational cost and intrinsically accelerating the exploration of conformational space, at the expense of introducing approximations that may affect both structural and kinetic properties^12,13^. Their accuracy must therefore be validated against experiment before they can be used predictively, particularly when the goal is not simply to reproduce binding for a single wild-type (WT) complex, but also to capture how binding is perturbed by mutations in either binding partner.

Multi-eGO is a hybrid transferable/structure-based atomistic-resolution model in which transferable prior interactions are re-weighted with contact probability information derived from reference data, such as structures, ensembles, and simulations^14,15^. The transferable prior potential provides a baseline description of generic interactions and conformational preferences, while the reference data introduce system-specific information through the observed contact probabilities. This framework has been shown to reproduce self-assembly processes^16,17^ and, in more recent applications, the binding thermodynamics of protein–ligand complexes^18^. In previous work, this approach quantitatively recovered the dissociation constants of small-molecule ligands binding to folded proteins, including T4-lysozyme and c-Src kinase as well as to disordered peptides like Aβ42^18^. This result was subsequently reproduced for trypsin^19^. Whether the same strategy, namely deriving the intermolecular interaction strength from nonspecific surface contacts without additional fitting to the bound complex, generalizes from small, relatively rigid ligands to flexible peptides remains unknown. It is also unclear whether the approach remains predictive when mutations are introduced in either the protein or the peptide. Allatom free-energy calculations can estimate the effects of mutations on PDZ–peptide binding^20,21^ and of protein mutations on binding in general^22^, but at a considerable computational cost for each mutant.

Here, we address both questions using the PDZ2 domain from protein tyrosine phosphatase basophil-like, PTP-BL, in complex with the peptide EQVTAV, a system for which high-resolution structural data and experimentally measured binding kinetics and equilibrium constants are available. Such data have been reported for the WT complex as well as for multiple PDZ2 point mutants, including L18A, L25A, T35G, V44A, and A53G, and for the same variants in complex with the peptide EQVSAV^23,24^, cf. Figure 1. In these experiments EQVSAV is the reference peptide^24^; here the model is trained on EQVTAV, so that the peptide mutation (T_−2_S, the reverse of the experimental S_−2_T) only removes a methyl group, and both peptides are simulated with WT PDZ2 and the five mutants (twelve complexes). These mutations are all conservative substitutions. We first assessed whether multi-eGO, trained without any peptide-specific tuning beyond the protocol previously established for small molecules, could reproduce the structural dynamics of PDZ2 as well as the binding kinetics and thermodynamics of the WT complex. We then tested, without additional training for each mutant, whether the same model could predict both the direction and magnitude of the changes in binding induced by mutations in either binding partner. This represents a substantially more stringent test of transferability than reproducing the properties of a single WT complex.

**Figure 1.**
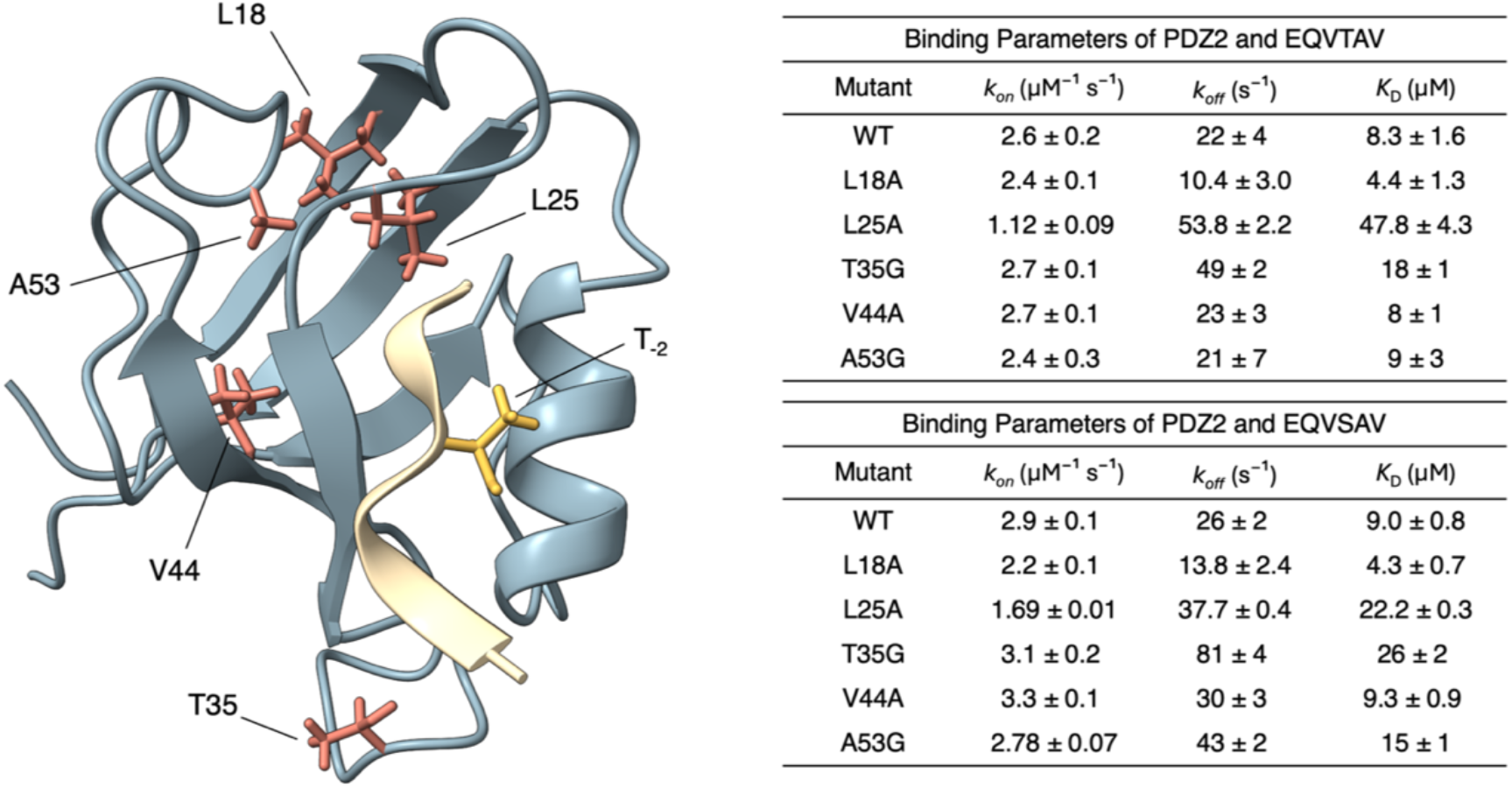
Structure and experimental binding data for the PDZ2 system. Left: cartoon representation of the PDZ2 structure (PDB: 1VJ6)^23^, with residues mutated in the present study shown as sticks. Right: experimentally measured association rate constants *k*_on_, dissociation rate constants k_off_, and equilibrium dissociation constants *K*_*D*_ for the wild-type and mutant PDZ2 systems in complex with EQVTAV (top) and EQVSAV (bottom), as reported in Ref.^24^. PDZ ligand residues are conventionally numbered relative to the C-terminus; for example, T_−2_ denotes the threonine residue located two positions upstream of the C-terminal residue.

Finally, we quantify how large the residual kinetic discrepancy is, by reweighting the simulated first-passage times a posteriori onto the experimental rate constants within a maximum-caliber framework. This separates a global acceleration, common to all complexes, from the system-specific kinetic errors of the model.

## METHODS

### Conventional Molecular Dynamics Simulations

All-atom, explicit-solvent reference simulations of the PDZ2 complex and of the apo systems (PDZ2 and EQVTAV) were performed using the GROMACS molecular dynamics (MD) software package^25^, employing the DES-Amber force field^26^ in combination with the TIP4P-D water model^27^. The starting structure of the protein complex was based on the NMR structure of the PDZ2 domain from PTP-BL, in complex with the C-terminal peptide of the adenomatous polyposis coli, APC protein (PDB: 1VJ6^23^), while the apo PDZ2 structure was based on the corresponding structure (PDB: 1GM1^28^). The peptide sequence was mutated to EQVTAV using the Mutate Residue tool in Maestro (Schrö-dinger, LLC, New York, NY^29^), and all systems were prepared and protonated using Maestro at pH 7.2, in agreement with the experimental conditions^24^.

Training simulations of the PDZ2 complex were performed by placing one peptide in the binding pocket and four additional peptides outside the binding pocket to sample interactions with the protein surface. Four independent replicas of the complex simulation, each 1 μs in length, were performed. All systems were solvated in a dodecahedral box in the presence of Na^+^ and Cl^−^ ions at a concentration of 800 mM, both to match the Na^+^ concentration of the experiments of Gianni et al.^24^ (0.4 M Na_2_SO_4_, 50 mM phosphate) and to neutralize the total charge of the system. The experiments used a Y43W pseudo-wild-type minimization in three sequential stages: steepest descent until the maximum force converged to less than 1000 kJ mol^−1^ nm^−1^, conjugate gradient until the maximum force converged to less than 100 kJ mol^−1^ nm^−1^, and L-BFGS until the maximum force converged to less than 10 kJ mol^−1^ nm^−1^. The minimized configurations were then equilibrated for 0.5 ns in the NPT ensemble at 283 K and 1 bar, with protein atoms restrained to their energy-minimized positions.

The simulations were integrated using the leap-frog algorithm with a 2-fs time step, and LINCS^30^ was used to constrain bonds involving hydrogen atoms. The cutoff for nonbonded interactions was set to 1 nm, and particle mesh Ewald^31^ was used to treat long-range electrostatic interactions. The temperature was maintained at 283 K using stochastic velocity rescaling^32^ with a coupling time constant of 1.0 ps, while the pressure was maintained at 1 bar using cell rescaling^33^ with a coupling time constant of 5.0 ps. A 1 μs NPT production run was then performed for each system under identical conditions. MD simulations of the apo systems were carried out following the same protocol described above. The temperature (283 K) follows earlier kinetic measurements on PDZ2^34^; the data of Ref.^24^ were measured at 277 K, a 2% difference in RT.

### Multi-eGO Molecular Dynamics Simulations

Multi-eGO is a hybrid transferable/structure-based atomistic force field in which all non-hydrogen atoms are explicitly represented, together with hydrogen atoms attached to the protein backbone^14,15^. Its functional form follows that of conventional molecular mechanics force fields, with Lennard-Jones (LJ) interactions used to describe all nonbonded interactions. Bonded terms, together with the default C^(12)^ LJ parameters, are transferable and derived from the GROMOS54A7 force field^35^. To improve the ability of multi-eGO to capture sequence-dependent changes in conformational properties, both dihedral and nonbonded prior interactions are reparametrized. Dihedral terms for all amino acids are optimized based on residue concentration as the training simulation, considering only the four unbound peptides.

All multi-eGO MD simulations were performed using stochastic dynamics integration with a 5-fs time step and a 25 ps relaxation time. The cutoff for LJ interactions was set to 2.5 nm. A 10% larger radius was used for the neighbor lists, which were updated every 20 steps. A range of *ε*_0_ values was tested to optimize the agreement between the training and multi-eGO simulations; each value of the intermolecular energy scale was evaluated with a 1 μs multi-eGO simulation, while 100 ns trajectories were used to evaluate the intramolecular energy scale. All simulations were performed using GROMACS 2024.

### Multi-eGO Modeling of Protein and Peptide Mutants

To further investigate protein–peptide binding kinetics and dissociation constants, five site-directed PDZ2 mutants (L18A, L25A, T35G, V44A, and A53G) were generated in addition to the wild-type PDZ2 complex. Mutant topologies and coordinate files were generated from the corresponding WT structure using the Gromologist package^44^. The corresponding intra- and inter-molecular contact matrices were derived from the WT matrices without repeating the computationally expensive all-atom PDZ2 and N-terminally dansylated peptides^24^; neither feature was modeled.

All systems were subjected to energy populations from the Coil Database and experimental helical propensities^36–39^. Nonbonded prior interactions are instead parameterized to reproduce experimental radii of gyration of intrinsically disordered proteins (IDPs), following approaches developed for CALVADOS^40^, Mpipi^41^, and other models^42,43^.

The nontransferable, structure-based component of the force field is derived from reference data, such as experimental structures, ensembles, and state-of-the-art simulations of the system of interest, and is reweighted relative to a prior simulation, which provides a simplified description of the same system. From the reference and prior simulations, pairwise atomic distances and the corresponding contact probabilities, 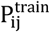 and 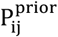, are extracted using the multi-eGO tools cmdata and make_mat.py. The atom-pair nonbonded interaction energy is then estimated by Bayesian reweighting as:

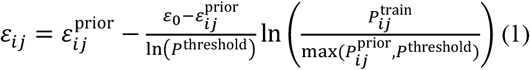

A detailed description of the model can be found in a previous work^15^, and the associated code and parameters are available on GitHub (https://github.com/multi-ego/multi-eGO) under the beta6 tag.

For the apo systems, a prior simulation was performed using the transferable multi-eGO prior, generated using only the transferable bonded and nonbonded interactions. For the PDZ2 complex, an intermolecular prior simulation was introduced, in which intramolecular interactions within each molecule are learned through the Bayesian procedure, while intermolecular interactions are described exclusively by transferable prior non-bonded interactions. The two sets of prior probabilities allow intra- and intermolecular interactions to be decoupled and assigned distinct energy scales (*ε*_0_). For intermolecular interactions, the prior simulation corresponds to the protein system at the same training step. A custom Python script removes all interactions involving the atoms deleted by each substitution, while all other interactions keep their WT values and energy scales. The backbone and Cβ atoms are thus retained for L→A and V→A, only the backbone for T→G and A→G, and for T→S the Cγ methyl group is removed while the hydroxyl oxygen is retained. Bonded terms, including the residue-specific dihedrals of the multi-eGO prior, are those of the new residue.

The same set of mutations was also generated in complex with the EQVSAV peptide, which was obtained from the EQVTAV peptide by a T→S substitution using the same approach described above.

### Calculation of Kinetic Parameters and Dissociation Constants

Following generation of the multi-eGO force field, binding and unbinding kinetics were characterized for each system. Binding times were obtained from 50 independent unbiased simulations, each 1 μs long at 283 K, containing the protein and four unbound peptides placed at random positions; a binding event was defined as the first of the four peptides reaching the bound state. Similarly, unbinding times were obtained from 50 independent simulations, each 4 μs long at 283 K, containing only the bound peptide and starting from the same conformation with random velocities extracted from a Max-well–Boltzmann distribution. The few trajectories that had not unbound within 4 μs were extended until unbinding occurred.

Single binding and unbinding times were determined using the COMMITTOR module of the PLUMED software package^45^. The COMMITTOR module was configured to stop each simulation once both the minimum protein–peptide distance (*d*_min_, calculated from the backbone hydrogen of L25 and the C-terminus backbone oxygens of the peptide) and the protein– peptide coordination number (*cn*, calculated using all the atoms of the protein and the peptide with a characteristic distance *R*_0_ of 0.3 nm) crossed predefined thresholds. The binding or unbinding time for each simulation was defined as the first time at which both *d*_min_ and *cn* crossed the corresponding thresholds. For the bound state, the thresholds were defined as 0 < *d*_min_ < 0.4 nm and 55 < *cn* < 150, while for the unbound state they were defined as 1.2 < *d*_min_ < 1000 nm and 0 < *cn* < 10. The bound-state thresholds were chosen from the distributions of the two variables in the bound basin of the reference simulations. Re-analysis of the stored time series with alternative thresholds does not change the statistical significance of the correlations for EQVTAV and EQVSAV (Tables S1 and S2).

The resulting distributions of binding and unbinding times were fitted to the exponential cumulative distribution function (CDF) expected for a Poisson process^46,47^ to estimate the mean binding and unbinding times, *τ*:

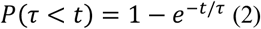

The Kolmogorov–Smirnov test^46^ was used to evaluate the goodness of fit. To obtain robust estimates of the associated uncertainties, a bootstrap resampling procedure was applied.

The kinetic parameters were calculated using the standard relationships:

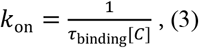

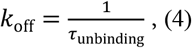

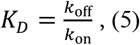

where [C] is the peptide concentration used in the simulations (0.0176 M, four peptides in a 378 nm^3^ box). The relation between K_D_ and the rate constants holds here because the binding and unbinding times are separated by more than three orders of magnitude^18,19^. The uncertainty in *K*_*D*_ was determined by analytical error propagation.

Pearson correlation coefficients, computed on ln K_D_, are reported in the Supporting Information with their 95% bootstrap confidence intervals, leave-one-out ranges, and one-sided p-values. The p-values were estimated with a Monte Carlo permutation test: the experimental and simulated values were perturbed with Gaussian noise according to their uncertainties, and the correlation of each perturbed dataset was compared with that obtained after randomly shuffling the simulated values (null hypothesis of no correlation).

### Comparison with NMR Data

Backbone ^3^J(H^N^–H^α^) couplings and ^1^D_NH_ residual dipolar couplings (RDCs) measured for PTP-BL PDZ2^28^ were back-calculated from the DES-Amber and multi-eGO trajectories: ^3^J couplings with the Karplus relation ^3^J = A cos^2^(φ − 60°) + B cos(φ − 60°) + C and three sets of parameters (A = 6.64, B = −1.43, C = 1.86 Hz^48^; A = 6.47, B = −1.43, C = 2.03 Hz^49^; A = 7.90, B = −1.05, C = 0.65 Hz^50^), taking the mean and the standard deviation over the three sets, and RDCs with PALES^51,52^ in SVD mode, fitting the alignment tensor to the experimental couplings frame by frame. Both were averaged over the trajectory, and the agreement is reported as RMSE (^3^J) and Q factor (RDCs)^52^.

### Maximum-Caliber Reweighting of the Kinetics

For each system and direction, we computed the acceleration factor A = k^sim^/k^exp^ from the mean first-passage times and fitted a single clock factor c across all systems and both directions, maximizing the smallest effective sample size. The residual factors f = A/c measure how much each simulated mean first-passage time must change to reproduce experiment (cf. Figure S1). The 50 first-passage times t_i_ were then reweighted onto an exponential distribution f_tgt_(t) (cf. Figures S2 and S3) with the experimental mean rescaled by c, using the minimum relative path entropy (maximum caliber) weights^53,54^ w_i_ ∝ f_tgt_(t_i_)/f_sim_(t_i_), with f_sim_ estimated by a gamma fit (cf. Figures S4 and S5). The cost of the correction was quantified by the Kish effective sample size^55^, N_eff_ = (Σ_i_ w_i_)^2^/Σ_i_ w_i_^2^. The derivation is given in the Supporting Information; calculations used the MaxCal package (https://github.com/multi-ego/MaxCal).

## RESULTS

The WT model is trained on explicit-solvent simulations of the complex with one bound and four free peptides. The contacts of the bound peptide enter the learned interactions, but the intermolecular energy scale is calibrated only on the free peptides, and neither affinity nor kinetics are used in training. Mutant models are derived from the WT model without further reference simulations or recalibration.

### Calibration of the intramolecular model

The reference data were obtained from explicit-solvent simulations of PDZ2 with one EQVTAV peptide bound and four additional peptides free in solution (Figure 2A). The bound peptide keeps its binding pose in all four replicas (peptide RMSD after fitting on PDZ2 below 0.6 nm and a stable hydrogen bond between the peptide C-terminal carboxylate and the backbone amide of L25; cf. Figure S6). This setup allowed us to sample the conformational dynamics of PDZ2 and the peptide while simultaneously characterizing nonspecific interactions between the peptide and the protein. Figure 2B shows the probability of contact formation between the peptide and the protein from the reference simulation, projected onto the PDZ2 surface.

**Figure 2.**
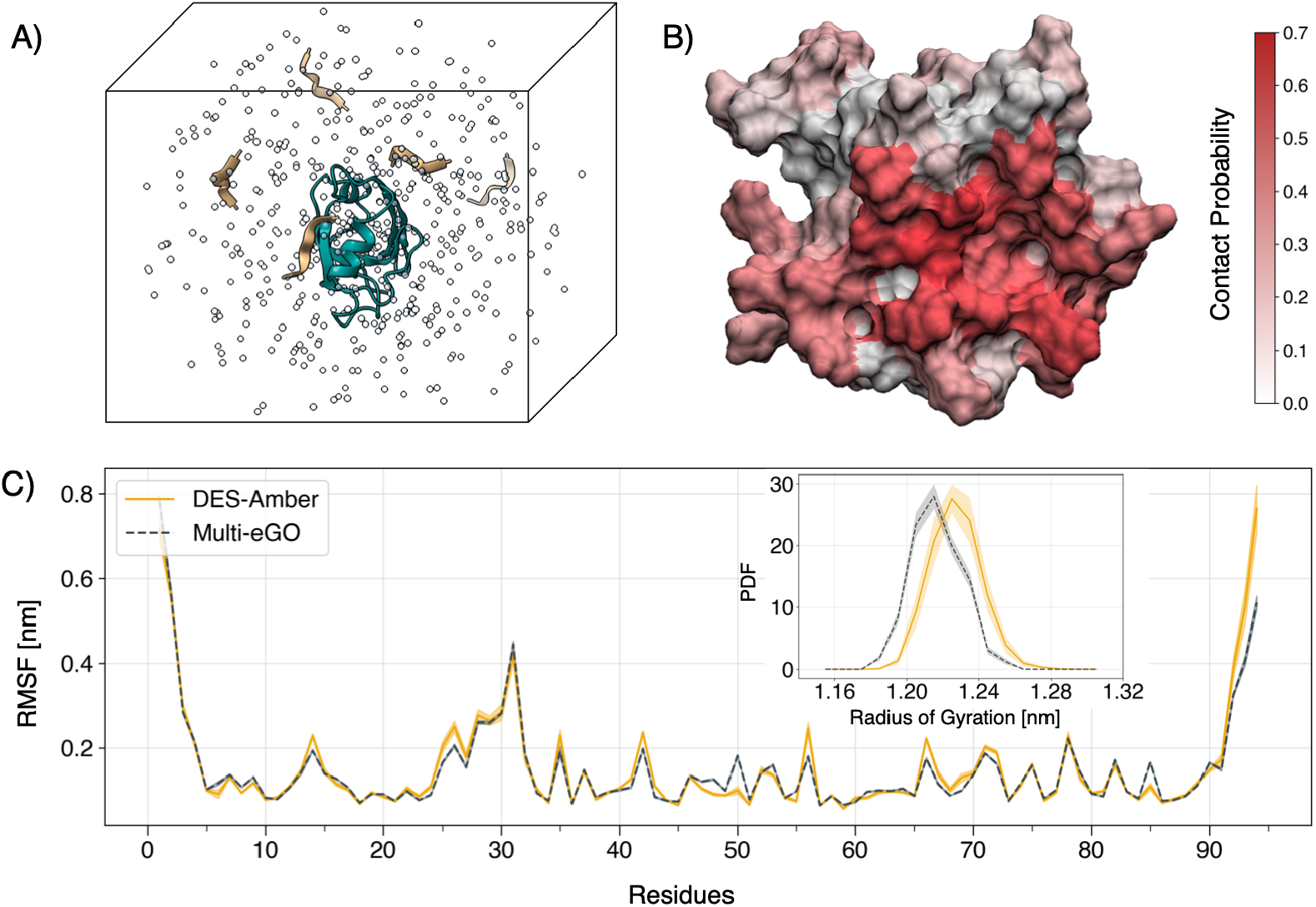
Reference simulations and structural properties of the PDZ2 system. (A) Representation of the simulated reference system, consisting of PDZ2 with one EQVTAV peptide bound and four additional EQVTAV peptides free in solution. (B) Surface representation of the per-residue intermolecular contact probability obtained from the PDZ2 reference simulation. (C) Comparison of the root-mean-square fluctuations (RMSF) obtained from the reference simulation using the DES-Amber force field and the multi-eGO simulation with the optimized 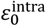 value of 0.28 kJ/mol. Inset: comparison of the probability density functions (PDFs) of the radius of gyration obtained from the DES-Amber reference simulation and the multi-eGO simulation with 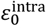 value of 0.28 kJ/mol. Shaded areas indicate uncertainties estimated by block analysis.

Each set of interactions is associated with an energy parameter (*ε*_0_) that must be determined. Intramolecular prior data were obtained from simulations of PDZ2 and the peptide using the transferable multi-eGO prior, which describes a molten-glob-ule-like ensemble (cf. Methods). For intramolecular interactions, 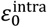 was set by minimizing the root-mean-square error (RMSE) between the root-mean-square fluctuations (RMSF) obtained from the multi-eGO and reference simulations. This procedure yielded an optimal value of 0.28 kJ/mol (cf. Figure S7A). With this choice of 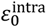, multi-eGO reproduces the equilibrium fluctuations of PDZ2 observed in the reference simulation (Figure 2C). The model also reproduces the probability distribution of the radius of gyration (Figure 2C, inset), which was not used in the fit, as well as the intramolecular contact map of the reference simulations (RMSE = 0.04, Figure S7B). Multi-eGO also reproduces the experimental ^3^J(H^N^–H^α^) and ^1^D_NH_ couplings of PDZ2^25^ with an accuracy close to that of the DES-Amber reference (^3^J RMSE 1.35 vs 1.23 Hz; RDC Q factor 0.60 vs 0.55; Figure S8).

### Multi-eGO predicts the wild-type binding affinity despite accelerated kinetics

Intermolecular prior data were obtained after setting the intramolecular energy scale 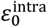 and retaining the transferable prior interactions only for intermolecular contacts. The energy scale for intermolecular interactions 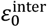 was determined, as previously introduced^18^, by minimizing the RMSE of the per-residue intermolecular contact probability, excluding the contacts formed by the bound peptide (Figure 3A,B). This procedure resulted in an optimal value of 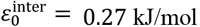.

**Figure 3.**
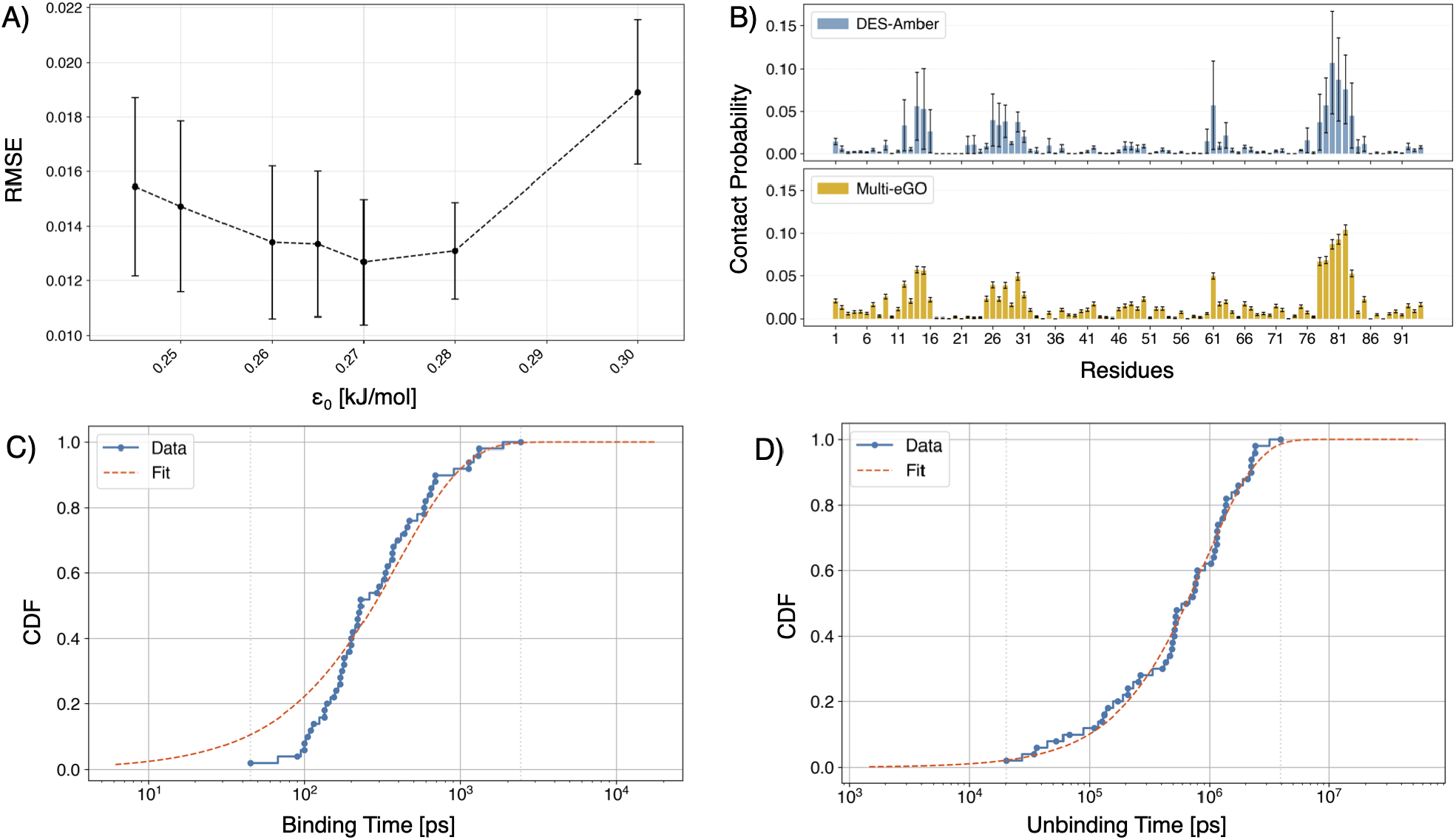
Optimization of the intermolecular energy scale and binding and unbinding kinetics of the PDZ2 system. (A) Root-mean-square error (RMSE) between the reference and multi-eGO intermolecular contact probabilities, excluding the bound EQVTAV peptide, as a function of 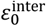 . Errors on the RMSE were obtained by first-order propagation of the block-averaging errors of the individual contact probabilities (maximum over block sizes), assuming independent errors. (B) Comparison of the intermolecular contact probabilities, excluding the bound EQVTAV peptide, obtained from the reference DES-Amber simulation and the production multi-eGO simulation performed with 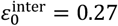 kJ/mol. Error bars were estimated by block analysis. (C) Cumulative distribution function (CDF) of the PDZ2 binding times (ps) obtained from 50 independent multi-eGO binding simulations, together with the corresponding exponential fit. (D) Cumulative distribution function (CDF) of the PDZ2 unbinding times (ps) obtained from 50 independent multi-eGO unbinding simulations, together with the corresponding exponential fit.

This approach is based on the hypothesis that the intermolecular energy scale is better captured by weak nonspecific surface interactions than by contacts at the native binding interface, whose contact probabilities are saturated by the high peptide concentration used in the reference simulation. At the same time, the prior model accounts for the translational and rotational entropy associated with the peptide concentration, allowing the concentration-dependent contribution from random clashes to be removed from the nonspecific interactions.

The binding of EQVTAV to PDZ2 was characterized by performing 50 independent binding and unbinding simulations and obtaining the CDFs of the binding and unbinding times (Figure 3C,D), which were fitted to an exponential distribution. The fit was good for the unbinding process (p = 0.91), whereas the binding times were less well described by an exponential (p = 0.06), consistent with the short lag time of a diffusive search (see below). The resulting mean binding and unbinding times were 391 ± 59 ps and 928,000 ± 182,000 ps, respectively, corresponding to estimated *k*_on_ and *k*_off_ values of 1.5 × 10^0^ ± 0.2 × 10^0^ μM^-1^ s^-1^ and 1.1 × 10^1^ ± 0.2 × 10^1^ s^-1^, respectively.

These kinetic rates differ by several orders of magnitude from the experimentally measured values (cf. Figure 1), with multi-eGO exhibiting substantially faster binding and unbinding kinetics, as expected from the simplified nature of the model. The simulated k_on_ (1.5 × 10^11^ M^−1^ s^−1^) is close to the diffusion limit of the model. Because of the low friction of the implicit-solvent Langevin dynamics (τ = 25 ps), diffusion in the model is about 25 times faster than in water, which raises the diffusion limit from about 4 × 10^9^ M^−1^ s^−1^ in water^2^ to about 1 × 10^11^ M^−1^ s^−1^. The experimental k_on_, instead, is three orders of magnitude below the diffusion limit in water. In the model, k_on_ therefore reflects the search time rather than the recognition step. The accelerated kinetics are, however, an important practical advantage of multi-eGO, as they allow extensive sampling of binding and unbinding processes within computationally accessible simulation times.

Despite the accelerated kinetics, the resulting equilibrium dissociation constant, K_D_ = 7.3 ± 1.4 μM, agrees with the experimental value (8.3 ± 1.6 μM) within the expected uncertainty. This result indicates that the procedure previously developed for small molecules can be transferred to protein–peptide interactions. Because the RMSE minimum in Figure 3A is shal-low, the absolute K_D_ carries an additional uncertainty from the choice of 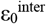, not included in the bootstrap errors, which largely cancels in relative affinities.

### Effects of protein and peptide mutations on binding affinity

A more stringent test of transferability is whether the same model can reproduce mutation-induced changes in binding without additional training. To model the effects of conservative mutations in PDZ2 and the peptide, we removed interactions involving atoms deleted by the mutation and updated the corresponding bonded interactions according to those of the mutated residue (cf. Methods and Figure 4).

**Figure 4.**
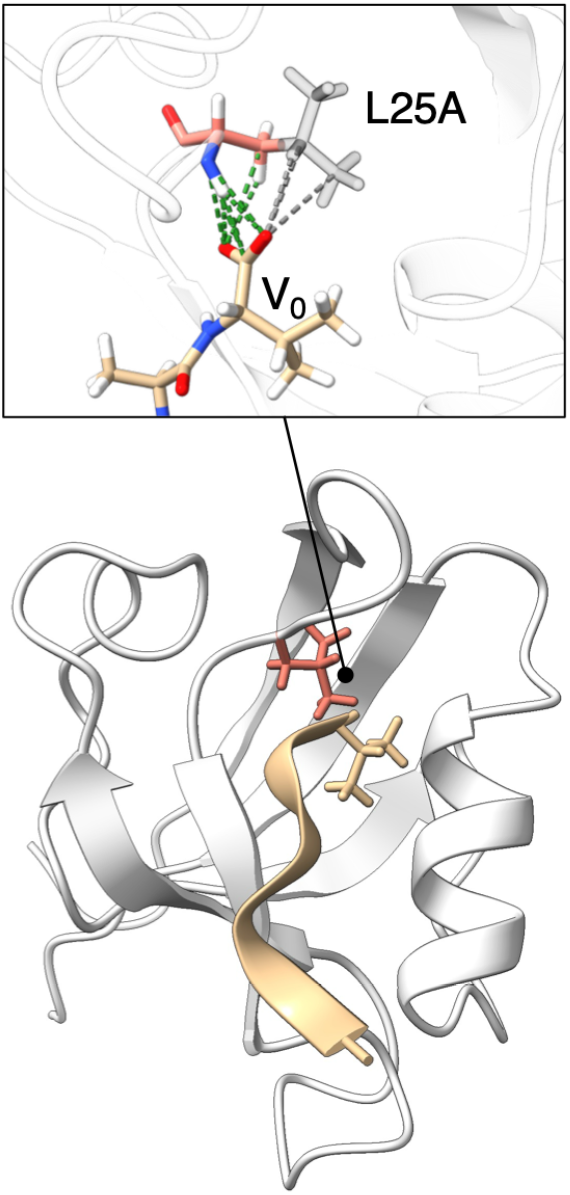
Schematic representation of the implementation of conservative mutations in multi-eGO, using the L25A mutation as a representative example. Inset: interactions between residue 25 and the peptide residue V_0_. Green dashed lines indicate backbone and Cβ contacts, which are retained after mutation; gray dashed lines indicate side-chain contacts involving atoms deleted by the L25A substitution, which are removed. The same procedure was applied to all mutations examined in this study.

After updating the simulation parameters for all mutations described in Figure 1, including five mutations in PDZ2 and one mutation in the peptide (EQVTAV→EQVSAV), we performed binding and unbinding simulations for each mutant system and calculated the corresponding *k*_on_ and *k*_off_ values (cf. Figures S9–S19 and Table 1). As observed for the WT system, the p-values associated with the fits of the unbinding time distributions were systematically higher than those for the binding time distributions. RMSD analyses in the bound state show that all mutants remain folded (Figure S20).

**Table 1.** Multi-eGO-predicted binding kinetics and equilibrium dissociation constants for WT and mutant PDZ2 in complex with EQVTAV and EQVSAV.

|  | EQVTAV |  |  | EQVSAV |  |  |
|---|---|---|---|---|---|---|
| Mu-tant | *k*<sub>*on*</sub> (10<sup>5</sup> µM<sup>−1</sup> s<sup>−1</sup>) | *k*<sub>*off*</sub> (10<sup>5</sup> s<sup>−1</sup>) | *K*<sub>D</sub> (µM) | *k*<sub>*on*</sub> (10<sup>5</sup> µM<sup>−1</sup> s<sup>−1</sup>) | *k*<sub>*off*</sub> (10<sup>5</sup> s<sup>−1</sup>) | *K*<sub>D</sub> (µM) |
| WT | 1.5±0.2 | 11.0±1.7 | 7.3±1.4 | 1.1±0.1 | 23.6±4.1 | 21±4 |
| L18A | 1.1±0.1 | 7.0±1.0 | 6.3±1.2 | 1.0±0.1 | 11.0±1.7 | 10±2 |
| L25A | 1.1±0.1 | 19.0±3.0 | 17±4 | 0.9±0.1 | 32.0±6.8 | 34±9 |
| T35G | 1.0±0.1 | 9.9±1.6 | 10.0±1.6 | 1.3±0.2 | 22.0±3.1 | 17±3 |
| V44A | 1.1±0.2 | 8.6±1.5 | 7.7±1.8 | 1.0±0.1 | 13.0±2.0 | 13.8±2.7 |
| A53G | 1.2±0.1 | 7.8±1.5 | 6.6±1.5 | 0.9±0.1 | 13.0±1.8 | 14.1±2.6 |

Consistent with the WT system, the simulated kinetics were several orders of magnitude faster than the experimental rates, whereas the equilibrium dissociation constants were on the same scale as the experimental values. A comparison with experimental measurements is shown in Figure 5. For the EQVTAV peptide, the *k*_23_ values showed poor correlation with the experimental values across the PDZ2 mutants, whereas the *k*_off_ values showed a stronger correlation. The predicted dissociation constants follow the experimental trend, which is dominated by L25A. In logarithmic scale the correlation remains high (Pearson r = 0.97 for log(K_D_); leave-one-out 0.89–0.99), and the ΔΔG error (mean absolute error, MAE, 0.97 kJ/mol) is lower than for a no-effect baseline (1.51 kJ/mol; Figure S21). The response is, however, compressed (slope ≈ 0.44), and the smallest effects (V44A and A53G, |ΔΔG| < 0.5 kJ/mol) are not resolved.

**Figure 5.**
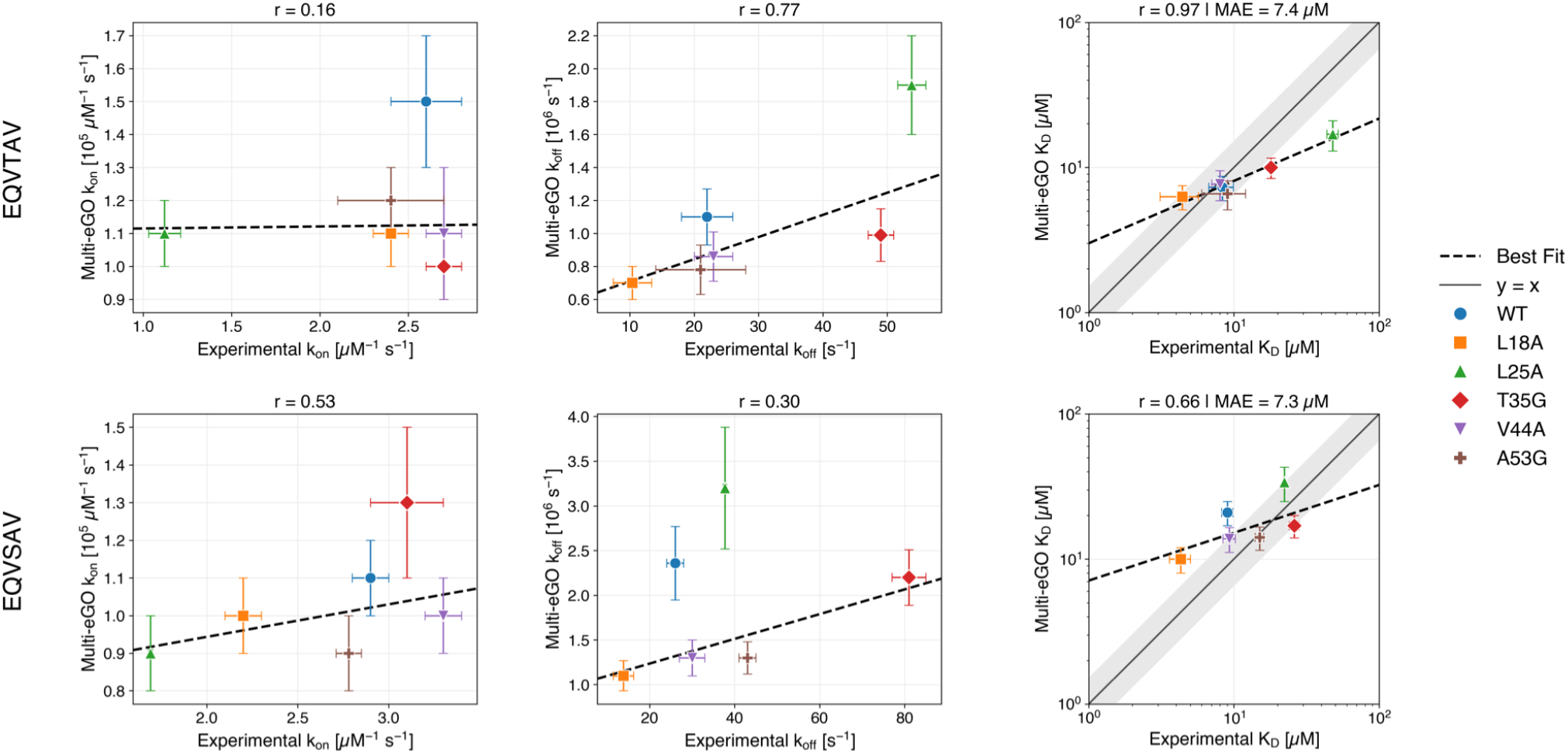
Correlation between multi-eGO-predicted and experimental association rate constants (k_on_), dissociation rate constants (k_off_), and equilibrium dissociation constants on logarithmic scale (K_D_) as a function of PDZ2 mutation, for (A) the wild-type peptide EQVTAV and (B) the peptide mutant EQVSAV. Points are colored by PDZ2 mutant, as indicated in the legend. Black dashed lines show the linear best fit. Correlation coefficients (r) are reported for all three quantities and mean absolute errors (MAE) are reported for K_D_. Error bars represent one standard deviation from bootstrap for multi-eGO and the experimental uncertainties reported in Ref. 21. In the logarithmic K_D_ plots, the shaded area marks deviations from the diagonal within ±1 kJ/mol.

When both binding partners were mutated, namely for the EQVSAV peptide in combination with the PDZ2 mutants, we observed only weak correlations for the individual kinetic rate constants. The dissociation constants are only moderately correlated with experiment in logarithmic scale (r = 0.66, p = 0.11), and the ΔΔG error (MAE 1.42 kJ/mol) is no better than the no-effect baseline (1.50 kJ/mol). The main error is the peptide sub-stitution itself: for the WT complex the model predicts ΔΔG = +2.5 kJ/mol for T_−2_S, against +0.2 kJ/mol in experiment. Accordingly, the double-mutant-cycle coupling energies between the PDZ2 mutations and the −2 position (−0.7 to −1.4 kJ/mol, with uncertainties of about 1 kJ/mol) do not resolve the only significant experimental coupling, that of L25A (−1.95 ± 0.53 kJ/mol). Pooling all twelve complexes gives a correlation of r = 0.61 (p = 0.04).

### A global clock factor accounts for most of the kinetic discrepancy

The comparison in Figure 5 conflates a global rescaling of time, which carries no information on the energetics, with system-specific errors. To separate them, we reweighted the simulated first-passage times onto the experimental rates within a maximum-caliber framework (cf. Methods). A single clock factor, c ≈ 6.3 × 10^4^, shared by all twelve complexes and both directions, accounts for most of the discrepancy. The residual factors f range from 0.35 to 1.65 (Figure 6), with a standard deviation of ln f of 0.37 (about 45%), and the effective sample sizes remain between 24 and 50 of the 50 trajectories. The measured rates are thus recovered by reweighting the simulated events, without any single trajectory dominating.

**Figure 6.**
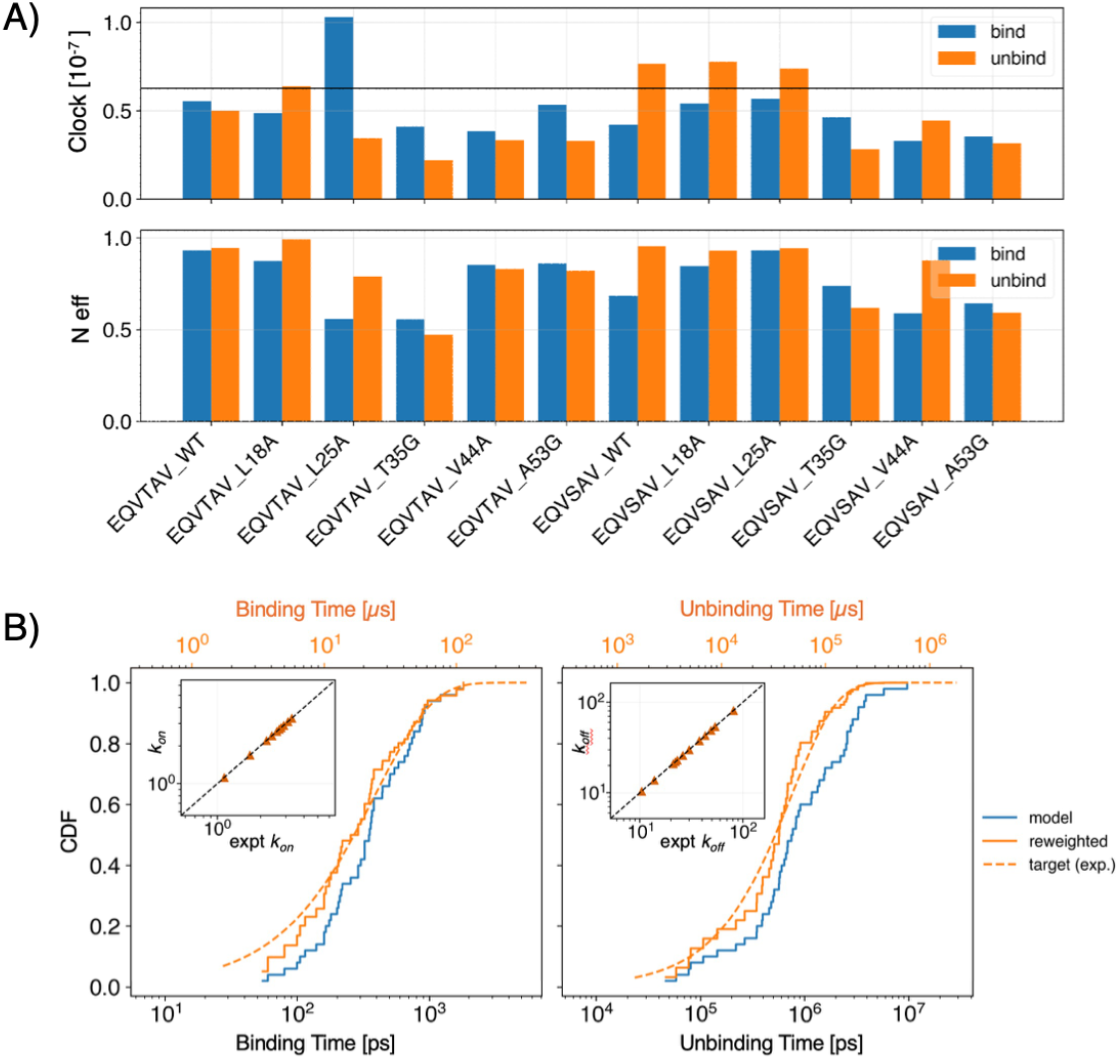
Maximum-caliber reweighting of the multi-eGO kinetics onto the experimental rate constants. (A) Top panel: system-specific clock factors, defined as the conversion from simulation to experimental time that would exactly reproduce the experimental binding (blue) and unbinding (orange) rate of each system. The horizontal line marks the single clock factor used for all systems (6.3 × 10^−8^ s ps^−1^). The system-specific values lie within a factor of about three of the common one; MaxCal reweighting corrects this residual discrepancy for each system. Bottom panel: normalized effective sample size (N_eff_) for binding and unbinding trajectories of all mutants after MaxCal reweighting. (B) CDFs of the binding (left) and unbinding (right) times of A53G:EQVTAV from the simulations, after reweighting, and for the exponential target (top axes: time rescaled by the clock factor). Insets: reweighted k_on_ and k_off_ against experimental rate constants for all systems.

The largest binding residual is that of L25A:EQVTAV (f_on_ = 1.65), the only mutant whose experimental k_on_ decreases markedly (from 2.6 to 1.12 μM^−1^ s^−1^), a recognition effect that a nearly diffusion-limited model cannot capture. For the other systems the experimental k_on_ varies little (2.2–3.3 μM^−1^ s^−1^), so their small binding residuals do not imply a correct description of recognition, and the changes in K_D_ arise mostly from k_off_. The residual errors in k_on_ and k_off_ partly cancel in K_D_, leaving a peptide-dependent offset (geometric mean of 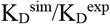 of 0.75 for EQVTAV and 1.40 for EQVSAV). Because the clock factor is fitted, this analysis does not yield absolute rate constants.

## DISCUSSION

We have shown that multi-eGO, trained with the protocol previously established for small-molecule ligands, reproduces the structural fluctuations of PDZ2 used for its calibration and predicts the WT binding affinity of the PDZ2 complex within the expected uncertainty. This extends the transferability of the underlying strategy, namely deriving the intermolecular interaction strength from nonspecific surface contacts rather than from the native binding interface, from small, rigid ligands to a flexible peptide. These results support the idea that the separation of entropic and enthalpic contributions achieved by this approach reflects general features of the model rather than arti-facts specific to small-molecule binding^18^.

The quality of the mutation predictions is uneven across observables, and this asymmetry is itself informative. Association rate constants *k*_on_ correlate poorly with experiment, dissociation rate constants *k*_off_ show a moderate correlation, whereas the equilibrium dissociation constants *K*_*D*_ show the best agreement. Because *K*_*D*_ is determined by the ratio 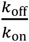, this pattern is consistent with correlated errors in the two rate constants partially canceling in their ratio, rather than with the model independently reproducing both kinetic rates. The acceleration plausibly reflects the smoother free-energy landscape of the simplified model, with lower effective barriers^55^. The maximum-caliber analysis makes this quantitative: once a single acceleration factor is removed, the residual errors are about 45%, are recovered by reweighting the simulated events, and partly cancel in K_D_.

For absolute values of *K*_D_, the bias depends on the peptide (underestimation for EQVTAV, overestimation for EQVSAV) and thus cannot be removed by a single global calibration. Relative affinities are better captured, but only for the largest effects of PDZ2 mutations (L25A, L18A) and with a response compressed about twofold. The weaker performance for the peptide mutation has a specific origin: T_−2_S removes the learned contacts of the threonine Cγ methyl group, and nothing in the model compensates for this loss, whereas PDZ2 discriminates poorly between serine and threonine at this position^21^, possibly because of differences in desolvation or local rearrangements of the pocket. This error propagates into the double-mutant cycles and is the main limitation of introducing peptide mutations without retraining.

It is also important to note that all mutations examined here are conservative substitutions that primarily remove or modify side-chain contacts without introducing new chemical functionality or changing the overall electrostatic character of the system. In the current implementation, the remaining contacts keep their WT strength, and the learned network keeps the WT fold, so that changes in dynamics and conformational entropy are captured only through the removed interactions and, for glycine, through the updated dihedral terms. Removing a side chain also removes its learned repulsive interactions, which can partly offset the loss of attractive ones. Whether this approach extends to non-conservative substitutions, such as changes in charge, proline substitutions that alter backbone flexibility, or bulkier side chains that introduce new steric contacts rather than simply removing existing ones, remains to be tested.

The ΔΔG error for conservative PDZ2 mutations with EQVTAV (about 1 kJ/mol) is within the typical error of allatom free-energy calculations^20–22^. Each mutant requires about 490 GPU-hours for 50 binding and 50 unbinding simulations, which give both rate constants and K_D_. For comparison, the allatom reference simulations used to train the WT model required 480 GPU-hours and contain no binding or unbinding event, as expected for experimental residence times of tens of milliseconds; they give access to neither the kinetics nor the affinity.

## CONCLUSIONS

We tested whether multi-eGO, previously validated for protein–small molecule binding, is transferable to protein–peptide interactions and can predict the effects of mutations on binding. Using the PDZ2 complex as a benchmark, we showed that multi-eGO, trained with the protocol established for small molecules, predicts the WT binding affinity within the expected uncertainty. We further showed that, without repeating the computationally expensive training step, the same model captures the direction and approximate size of the largest effects of conservative PDZ2 mutations on the equilibrium dissociation constant, with a compressed response, whereas smaller effects, the peptide mutation and the coupling between peptide and protein mutations are not resolved, and the individual rate constants agree less well with experiment.

A maximum-caliber reweighting shows that a single acceleration factor accounts for most of the kinetic discrepancy, and that the remaining system-specific errors are recovered by re-weighting the simulated events: up to a global time scale, the sampled binding and unbinding events are consistent with the experimental kinetics. Taken together, these results support the use of multi-eGO as a computationally efficient framework for studying protein recognition mechanisms and for estimating the effects of conservative protein mutations on binding affinity. At the same time, absolute kinetic rates, peptide mutations and non-conservative substitutions remain important directions for future work.

## ASSOCIATED CONTENT

### Supporting Information

Optimization of the intramolecular energy scale; binding and unbinding time distributions for all systems; correlation analysis in free-energy units and double-mutant-cycle coupling energies; intramolecular contact maps; stability of the bound pose in the reference simulations; comparison with NMR data; structural properties of the mutants; robustness to the state thresholds; derivation and additional results of the maximum-caliber reweighting (PDF).

### Author Contributions

CA performed and analyzed all the simulations. BS contributed to the setup and analysis of all the simulations. FBT worked on a proof of concept of the implementation of the mutations. BS, FBT, RC, and CC developed the multi-eGO model. SG, RC, and CC designed the project. RC and CC supervised the work. CA, BS, and CC wrote the manuscript with contributions of all authors.

## ACKNOWLEDGMENT

The authors acknowledge CINECA for an award under the ISCRA initiative, for the availability of high-performance computing resources and support.

## ABBREVIATIONS

MD: molecular dynamics
WT: wild type
LJ: Lennard-Jones
RMSF: root-mean square fluctuations
RMSE: root-mean square error
CDF: cumulative distribution function
PDF: probability density function
PDZ: PSD-95/Discs-large/ZO-1
PTP-BL: protein tyrosine phosphatase basophil-like
MAE: mean absolute error
MaxCal: maximum caliber
KS: Kolmogorov–Smirnov
RDC: residual dipolar coupling
SVD: singular value decomposition

## DATA AND SOFTWARE AVAILABILITY STATEMENT

Simulation data are publicly available on Zenodo (DOI: 10.5281/zenodo.22691022); the multi-eGO code and parameters are publicly available on GitHub at https://github.com/multi-ego/multi-eGO, use the beta6 tag for a snapshot of the repository associated to this paper. The maximum-caliber reweighting was performed with the MaxCal package (https://github.com/multi-ego/MaxCal); first-passage times, collective-variable time series, reweighting weights and comparison with NMR data are included in the Zenodo record.

