## Supporting information for "Assessing multi-eGO Predictions of PDZ2–Peptide Binding across Mutations"

### xFG SUPPORTING INFORMATION

**Supporting Text: Maximum-Caliber Reweighting of the First-Passage Times**

**Time-scale calibration.** For each system $s$ and direction $d$ (binding or unbinding), the acceleration factor is $A_{s,d} = k_{s,d}^{\mathrm{sim}}/k_{s,d}^{\mathrm{exp}}$, where the rate constants are obtained from the mean first-passage times and binding is treated as a pseudo-first-order process at the peptide concentration of the simulations ([C] = 0.0176 M). A single clock factor $c$ is fitted across the twelve systems and both directions, which leaves the residual factors $f_{s,d} = A_{s,d}/c$, that is, the factors by which each simulated mean first-passage time must change to reproduce experiment. Because $c$ cancels in every ratio of rate constants, only the residual factors carry information on the model. We used the value of $c$ that maximizes the smallest effective sample size among the 24 reweightings (balanced clock, $c \approx 6.3 \times 10^4$; see below and Figure S1). The geometric mean of the acceleration factors is a natural alternative, but it would require the mean binding time of L25A:EQVTAV to be lengthened by more than a factor of two, which is beyond what reweighting can achieve (Figure S1A).

**Maximum-caliber weights.** Let $P(\omega)$ be the probability of a trajectory $\omega$ in the multi-eGO ensemble and $t(\omega)$ its first-passage time. The distribution of the simulated first-passage times is

$$f_{\mathrm{sim}}(t) = \int \mathrm{d}\omega\, P(\omega)\, \delta(t - t(\omega)) \tag{S1}$$

Experiment provides a rate constant for each system and direction, which we represent as a Poisson process on the time scale of the model, $f_{\mathrm{tgt}}(t) = \tau_{\mathrm{tgt}}^{-1} \exp(-t/\tau_{\mathrm{tgt}})$, with $\tau_{\mathrm{tgt}} = 1/(c\, k_{\mathrm{off}}^{\mathrm{exp}})$ for unbinding and $\tau_{\mathrm{tgt}} = 1/(c\, [\mathrm{C}]\, k_{\mathrm{on}}^{\mathrm{exp}})$ for binding. Among all path ensembles $P'(\omega)$ whose distribution of first-passage times equals $f_{\mathrm{tgt}}$, the maximum-caliber principle (refs 53 and 54 of the main text) selects the one that is closest to the simulated ensemble in relative entropy,

$$\min_{P'} \int \mathrm{d}\omega\, P'(\omega) \ln \frac{P'(\omega)}{P(\omega)} \tag{S2}$$

subject to the constraint

$$\int \mathrm{d}\omega\, P'(\omega)\, \delta(t - t(\omega)) = f_{\mathrm{tgt}}(t) \qquad \text{for all } t \tag{S3}$$

which also ensures normalization. Introducing a Lagrange multiplier $\mu(t)$ for each value of $t$ and setting to zero the functional derivative of the Lagrangian with respect to $P'(\omega)$ gives

$$P'(\omega) = P(\omega) \exp[-1 - \mu(t(\omega))] \tag{S4}$$

Integrating eq S4 over all trajectories with first-passage time $t$ and imposing eq S3 fixes $\exp[-1 - \mu(t)] = f_{\mathrm{tgt}}(t)/f_{\mathrm{sim}}(t)$, so that

$$P'(\omega) = P(\omega)\, \frac{f_{\mathrm{tgt}}(t(\omega))}{f_{\mathrm{sim}}(t(\omega))} \tag{S5}$$

The minimum-perturbation solution is thus the importance ratio between the target and the simulated distributions of first-passage times. Because the correction depends on $\omega$ only through $t(\omega)$, the conditional ensemble at fixed first-passage time is unchanged, $P'(\omega|t) = P(\omega|t)$: the reweighting changes how much each trajectory contributes, not what it contains. For the $N = 50$ first-passage times $t_i$ of each system and direction, the weights are

$$w_i \propto \frac{f_{\mathrm{tgt}}(t_i)}{f_{\mathrm{sim}}(t_i)}, \qquad \sum_i w_i = 1 \tag{S6}$$

and the average of any per-trajectory observable $O$ that is consistent with the experimental kinetics is $\langle O \rangle' = \Sigma_i\, w_i O_i$. The density $f_{\mathrm{sim}}$ was estimated with a gamma distribution fitted to the simulated times by maximum likelihood (Figures S4 and S5). The distributions before and after reweighting are compared with the target in Figures S2 and S3.

**Cost of the reweighting.** The statistical cost of the correction is measured by the Kish effective sample size (ref 55 of the main text), $N_{\mathrm{eff}} = (\Sigma_i\, w_i)^2/\Sigma_i\, w_i^2$. When both the simulated and the target distributions are exponential, with $\tau_{\mathrm{tgt}} = f\,\tau_{\mathrm{sim}}$, the weights are $w(t) \propto \exp(-\theta t)$ with $\theta\tau_{\mathrm{sim}} = 1/f - 1$, and

$$\frac{N_{\mathrm{eff}}}{N} = \frac{\langle w\rangle^2}{\langle w^2\rangle} = 2f - f^2 \tag{S7}$$

where the averages are taken over the simulated distribution. The result does not depend on $N$. Shortening the mean first-passage time is therefore inexpensive (75% of the sample is retained for $f = 0.5$), whereas lengthening it becomes impossible as $f$ approaches 2, where $\langle w^2\rangle$ diverges. Eq S7 is compared in Figure S1B with the effective sample sizes obtained from the simulated times. All calculations were performed with the maxcal-target tool of the MaxCal package (https://github.com/multi-ego/MaxCal).

#### Supporting Figures:

| System | Thresholds | kon ($10^5$ μM$^{-1}$ s$^{-1}$) | koff ($10^5$ s$^{-1}$) | KD (μM) |
|---|---|---|---|---|
| **EQVTAV WT** | Original | 1.47 ± 0.18 | 10.7 ± 1.7 | 7.3 ± 1.5 |
| | Bound, checked every 1 ps | 1.52 ± 0.20 | 10.7 ± 1.7 | 7.0 ± 1.4 |
| | Bound d < 0.45 nm | 1.47 ± 0.18 | 10.7 ± 1.7 | 7.3 ± 1.5 |
| | Bound d < 0.5 nm | 1.47 ± 0.18 | 10.7 ± 1.7 | 7.3 ± 1.5 |
| | Bound c > 45 | 1.53 ± 0.20 | 10.7 ± 1.7 | 7.0 ± 1.4 |
| | Unbound d > 1.0 nm | 1.47 ± 0.18 | 11.8 ± 1.6 | 8.1 ± 1.5 |
| | Unbound d > 1.1 nm | 1.47 ± 0.18 | 11.3 ± 1.7 | 7.7 ± 1.5 |
| | Unbound c < 12 | 1.47 ± 0.18 | 11.2 ± 1.7 | 7.7 ± 1.5 |
| | Unbound c < 15 | 1.47 ± 0.18 | 15.3 ± 2.6 | 10.5 ± 2.2 |

*Table S1a. Kinetic parameters obtained with the original COMMITTOR thresholds and with alternative thresholds, varied one at a time while keeping all other thresholds at their original values. Since each simulation stops when the original criterion is met, only less strict thresholds could be tested. In the original analysis, the bound-state condition was checked every 5 ps, as in the COMMITTOR; "Bound, checked every 1 ps" uses instead the first frame at which the condition is met. Errors are bootstrap standard deviations. The "Original" values were recomputed with the same procedure used for the alternative thresholds and may differ slightly from those reported in the main text because of the bootstrap resampling.*

| System | Thresholds | kon ($10^5$ μM$^{-1}$ s$^{-1}$) | koff ($10^5$ s$^{-1}$) | KD (μM) |
|---|---|---|---|---|
| **EQVTAV L18A** | Original | 1.11 ± 0.14 | 7.0 ± 1.0 | 6.3 ± 1.2 |
| | Bound, checked every 1 ps | 1.14 ± 0.15 | 7.0 ± 1.0 | 6.1 ± 1.2 |
| | Bound d < 0.45 nm | 1.11 ± 0.14 | 7.0 ± 1.0 | 6.3 ± 1.2 |
| | Bound d < 0.5 nm | 1.11 ± 0.14 | 7.0 ± 1.0 | 6.3 ± 1.2 |
| | Bound c > 45 | 1.16 ± 0.15 | 7.0 ± 1.0 | 6.0 ± 1.2 |
| | Unbound d > 1.0 nm | 1.11 ± 0.14 | 7.2 ± 1.1 | 6.5 ± 1.3 |
| | Unbound d > 1.1 nm | 1.11 ± 0.14 | 7.2 ± 1.1 | 6.5 ± 1.3 |
| | Unbound c < 12 | 1.11 ± 0.14 | 7.8 ± 1.2 | 7.0 ± 1.4 |
| | Unbound c < 15 | 1.11 ± 0.14 | 9.6 ± 1.5 | 8.7 ± 1.7 |

*Table S2b. Kinetic parameters of EQVTAV L18A with the original and alternative COMMITTOR thresholds (see Table S1a for details).*

| System | Thresholds | kon ($10^5$ μM$^{-1}$ s$^{-1}$) | koff ($10^5$ s$^{-1}$) | KD (μM) |
|---|---|---|---|---|
| **EQVTAV L25A** | Original | 1.09 ± 0.14 | 18.7 ± 3.1 | 17.1 ± 3.6 |
| | Bound, checked every 1 ps | 1.17 ± 0.16 | 18.7 ± 3.1 | 16.0 ± 3.4 |
| | Bound d < 0.45 nm | 1.09 ± 0.14 | 18.7 ± 3.1 | 17.1 ± 3.6 |
| | Bound d < 0.5 nm | 1.09 ± 0.14 | 18.7 ± 3.1 | 17.1 ± 3.6 |
| | Bound c > 45 | 1.15 ± 0.16 | 18.7 ± 3.1 | 16.3 ± 3.5 |
| | Unbound d > 1.0 nm | 1.09 ± 0.14 | 21.9 ± 3.5 | 20.0 ± 4.1 |
| | Unbound d > 1.1 nm | 1.09 ± 0.14 | 20.6 ± 3.4 | 18.9 ± 4.0 |
| | Unbound c < 12 | 1.09 ± 0.14 | 20.7 ± 3.1 | 19.0 ± 3.8 |
| | Unbound c < 15 | 1.09 ± 0.14 | 23.5 ± 3.6 | 21.6 ± 4.4 |

*Table S3c. Kinetic parameters of EQVTAV L25A with the original and alternative COMMITTOR thresholds (see Table S1a for details).*

| System | Thresholds | kon ($10^5$ μM$^{-1}$ s$^{-1}$) | koff ($10^5$ s$^{-1}$) | KD (μM) |
|---|---|---|---|---|
| **EQVTAV T35G** | Original | 0.97 ± 0.10 | 9.7 ± 1.3 | 10.1 ± 1.7 |
| | Bound, checked every 1 ps | 1.03 ± 0.10 | 9.7 ± 1.3 | 9.4 ± 1.6 |
| | Bound d < 0.45 nm | 0.98 ± 0.10 | 9.7 ± 1.3 | 10.0 ± 1.7 |
| | Bound d < 0.5 nm | 0.98 ± 0.10 | 9.7 ± 1.3 | 10.0 ± 1.7 |
| | Bound c > 45 | 1.03 ± 0.10 | 9.7 ± 1.3 | 9.5 ± 1.6 |
| | Unbound d > 1.0 nm | 0.97 ± 0.10 | 11.2 ± 1.4 | 11.6 ± 1.9 |
| | Unbound d > 1.1 nm | 0.97 ± 0.10 | 10.1 ± 1.4 | 10.4 ± 1.8 |
| | Unbound c < 12 | 0.97 ± 0.10 | 10.7 ± 1.5 | 11.1 ± 1.9 |
| | Unbound c < 15 | 0.97 ± 0.10 | 13.0 ± 1.5 | 13.4 ± 2.1 |

*Table S4d. Kinetic parameters of EQVTAV T35G with the original and alternative COMMITTOR thresholds (see Table S1a for details).*

| System | Thresholds | kon ($10^5$ μM$^{-1}$ s$^{-1}$) | koff ($10^5$ s$^{-1}$) | KD (μM) |
|---|---|---|---|---|
| **EQVTAV V44A** | Original | 1.10 ± 0.18 | 8.7 ± 1.6 | 7.9 ± 1.9 |
| | Bound, checked every 1 ps | 1.14 ± 0.19 | 8.7 ± 1.6 | 7.6 ± 1.9 |
| | Bound d < 0.45 nm | 1.10 ± 0.18 | 8.7 ± 1.6 | 7.9 ± 1.9 |
| | Bound d < 0.5 nm | 1.10 ± 0.18 | 8.7 ± 1.6 | 7.9 ± 1.9 |
| | Bound c > 45 | 1.13 ± 0.18 | 8.7 ± 1.6 | 7.7 ± 1.9 |
| | Unbound d > 1.0 nm | 1.10 ± 0.18 | 10.3 ± 2.0 | 9.3 ± 2.4 |
| | Unbound d > 1.1 nm | 1.10 ± 0.18 | 9.1 ± 1.6 | 8.2 ± 2.0 |
| | Unbound c < 12 | 1.10 ± 0.18 | 9.5 ± 1.8 | 8.6 ± 2.1 |
| | Unbound c < 15 | 1.10 ± 0.18 | 11.3 ± 2.1 | 10.3 ± 2.5 |

*Table S5f. Kinetic parameters of EQVTAV V44A with the original and alternative COMMITTOR thresholds (see Table S1a for details).*

| System | Thresholds | kon ($10^5$ μM$^{-1}$ s$^{-1}$) | koff ($10^5$ s$^{-1}$) | KD (μM) |
|---|---|---|---|---|
| **EQVTAV A53G** | Original | 1.18 ± 0.14 | 7.9 ± 1.4 | 6.7 ± 1.4 |
| | Bound, checked every 1 ps | 1.23 ± 0.15 | 7.9 ± 1.4 | 6.4 ± 1.4 |
| | Bound d < 0.45 nm | 1.18 ± 0.14 | 7.9 ± 1.4 | 6.7 ± 1.4 |
| | Bound d < 0.5 nm | 1.18 ± 0.14 | 7.9 ± 1.4 | 6.7 ± 1.4 |
| | Bound c > 45 | 1.24 ± 0.15 | 7.9 ± 1.4 | 6.4 ± 1.3 |
| | Unbound d > 1.0 nm | 1.18 ± 0.14 | 8.2 ± 1.5 | 6.9 ± 1.5 |
| | Unbound d > 1.1 nm | 1.18 ± 0.14 | 8.1 ± 1.4 | 6.8 ± 1.4 |
| | Unbound c < 12 | 1.18 ± 0.14 | 8.9 ± 1.3 | 7.5 ± 1.4 |
| | Unbound c < 15 | 1.18 ± 0.14 | 10.5 ± 1.5 | 8.8 ± 1.6 |

*Table S6g. Kinetic parameters of EQVTAV A53G with the original and alternative COMMITTOR thresholds (see Table S1a for details).*

| System | Thresholds | kon ($10^5$ μM$^{-1}$ s$^{-1}$) | koff ($10^5$ s$^{-1}$) | KD (μM) |
|---|---|---|---|---|
| **EQVSAV WT** | Original | 1.10 ± 0.11 | 23.6 ± 4.2 | 21.5 ± 4.3 |
| | Bound, checked every 1 ps | 1.14 ± 0.12 | 23.6 ± 4.2 | 20.8 ± 4.3 |
| | Bound d < 0.45 nm | 1.10 ± 0.11 | 23.6 ± 4.2 | 21.5 ± 4.3 |
| | Bound d < 0.5 nm | 1.10 ± 0.11 | 23.6 ± 4.2 | 21.5 ± 4.3 |
| | Bound c > 45 | 1.17 ± 0.12 | 23.6 ± 4.2 | 20.2 ± 4.2 |
| | Unbound d > 1.0 nm | 1.10 ± 0.11 | 25.9 ± 4.2 | 23.6 ± 4.5 |
| | Unbound d > 1.1 nm | 1.10 ± 0.11 | 25.3 ± 4.3 | 23.0 ± 4.6 |
| | Unbound c < 12 | 1.10 ± 0.11 | 25.5 ± 4.3 | 23.2 ± 4.6 |
| | Unbound c < 15 | 1.10 ± 0.11 | 29.2 ± 4.9 | 26.6 ± 5.2 |

*Table S7h. Kinetic parameters of EQVSAV WT with the original and alternative COMMITTOR thresholds (see Table S1a for details).*

| System | Thresholds | kon ($10^5$ μM$^{-1}$ s$^{-1}$) | koff ($10^5$ s$^{-1}$) | KD (μM) |
|---|---|---|---|---|
| **EQVSAV L18A** | Original | 1.04 ± 0.11 | 10.6 ± 1.7 | 10.2 ± 2.0 |
| | Bound, checked every 1 ps | 1.07 ± 0.12 | 10.6 ± 1.7 | 9.9 ± 2.0 |
| | Bound d < 0.45 nm | 1.04 ± 0.11 | 10.6 ± 1.7 | 10.2 ± 2.0 |
| | Bound d < 0.5 nm | 1.04 ± 0.11 | 10.6 ± 1.7 | 10.2 ± 2.0 |
| | Bound c > 45 | 1.13 ± 0.13 | 10.6 ± 1.7 | 9.4 ± 1.9 |
| | Unbound d > 1.0 nm | 1.04 ± 0.11 | 13.1 ± 2.1 | 12.6 ± 2.4 |
| | Unbound d > 1.1 nm | 1.04 ± 0.11 | 11.3 ± 1.8 | 10.9 ± 2.1 |
| | Unbound c < 12 | 1.04 ± 0.11 | 13.9 ± 2.4 | 13.3 ± 2.8 |
| | Unbound c < 15 | 1.04 ± 0.11 | 16.9 ± 2.9 | 16.2 ± 3.3 |

*Table S8i. Kinetic parameters of EQVSAV L18A with the original and alternative COMMITTOR thresholds (see Table S1a for details).*

| System | Thresholds | kon ($10^5$ µM$^{-1}$ s$^{-1}$) | koff ($10^5$ s$^{-1}$) | KD (µM) |
|---|---|---|---|---|
| **EQVSAV L25A** | Original | 0.93 ± 0.13 | 31.9 ± 6.7 | 34.3 ± 8.6 |
| | Bound, checked every 1 ps | 0.95 ± 0.13 | 31.9 ± 6.7 | 33.7 ± 8.4 |
| | Bound d < 0.45 nm | 0.95 ± 0.12 | 31.9 ± 6.7 | 33.5 ± 8.3 |
| | Bound d < 0.5 nm | 0.95 ± 0.12 | 31.9 ± 6.7 | 33.5 ± 8.3 |
| | Bound c > 45 | 0.99 ± 0.13 | 31.9 ± 6.7 | 32.4 ± 8.0 |
| | Unbound d > 1.0 nm | 0.93 ± 0.13 | 40.5 ± 7.3 | 43.5 ± 9.9 |
| | Unbound d > 1.1 nm | 0.93 ± 0.13 | 34.3 ± 6.7 | 36.9 ± 8.8 |
| | Unbound c < 12 | 0.93 ± 0.13 | 41.1 ± 8.0 | 44.3 ± 10.5 |
| | Unbound c < 15 | 0.93 ± 0.13 | 53.4 ± 10.4 | 57.4 ± 13.7 |

*Table S9i. Kinetic parameters of EQVSAV L25A with the original and alternative COMMITTOR thresholds (see Table S1a for details).*

| System | Thresholds | kon ($10^5$ µM$^{-1}$ s$^{-1}$) | koff ($10^5$ s$^{-1}$) | KD (µM) |
|---|---|---|---|---|
| **EQVSAV T35G** | Original | 1.30 ± 0.16 | 22.0 ± 3.0 | 16.9 ± 3.1 |
| | Bound, checked every 1 ps | 1.38 ± 0.17 | 22.0 ± 3.0 | 15.9 ± 2.9 |
| | Bound d < 0.45 nm | 1.30 ± 0.16 | 22.0 ± 3.0 | 16.9 ± 3.1 |
| | Bound d < 0.5 nm | 1.34 ± 0.16 | 22.0 ± 3.0 | 16.4 ± 3.0 |
| | Bound c > 45 | 1.43 ± 0.19 | 22.0 ± 3.0 | 15.4 ± 2.9 |
| | Unbound d > 1.0 nm | 1.30 ± 0.16 | 24.1 ± 3.0 | 18.6 ± 3.2 |
| | Unbound d > 1.1 nm | 1.30 ± 0.16 | 22.6 ± 3.1 | 17.4 ± 3.2 |
| | Unbound c < 12 | 1.30 ± 0.16 | 23.3 ± 3.3 | 18.0 ± 3.3 |
| | Unbound c < 15 | 1.30 ± 0.16 | 25.8 ± 3.9 | 19.9 ± 3.8 |

*Table S10l. Kinetic parameters of EQVSAV T35G with the original and alternative COMMITTOR thresholds (see Table S1a for details).*

| System | Thresholds | kon ($10^5$ µM$^{-1}$ s$^{-1}$) | koff ($10^5$ s$^{-1}$) | KD (µM) |
|---|---|---|---|---|
| **EQVSAV V44A** | Original | 0.98 ± 0.13 | 13.3 ± 1.9 | 13.6 ± 2.6 |
| | Bound, checked every 1 ps | 1.03 ± 0.14 | 13.3 ± 1.9 | 13.0 ± 2.6 |
| | Bound d < 0.45 nm | 0.98 ± 0.13 | 13.3 ± 1.9 | 13.6 ± 2.6 |
| | Bound d < 0.5 nm | 0.98 ± 0.13 | 13.3 ± 1.9 | 13.6 ± 2.6 |
| | Bound c > 45 | 1.09 ± 0.15 | 13.3 ± 1.9 | 12.2 ± 2.4 |
| | Unbound d > 1.0 nm | 0.98 ± 0.13 | 16.6 ± 2.6 | 16.9 ± 3.5 |
| | Unbound d > 1.1 nm | 0.98 ± 0.13 | 15.2 ± 2.2 | 15.5 ± 3.1 |
| | Unbound c < 12 | 0.98 ± 0.13 | 14.7 ± 2.1 | 15.0 ± 2.9 |
| | Unbound c < 15 | 0.98 ± 0.13 | 16.7 ± 2.4 | 17.1 ± 3.3 |

*Table S11m. Kinetic parameters of EQVSAV V44A with the original and alternative COMMITTOR thresholds (see Table S1a for details).*

| System | Thresholds | kon ($10^5$ μM$^{-1}$ s$^{-1}$) | koff ($10^5$ s$^{-1}$) | KD (μM) |
|---|---|---|---|---|
| **EQVSAV A53G** | Original | 0.91 ± 0.11 | 12.8 ± 1.8 | 14.0 ± 2.6 |
| | Bound, checked every 1 ps | 0.94 ± 0.12 | 12.8 ± 1.8 | 13.6 ± 2.5 |
| | Bound d < 0.45 nm | 0.91 ± 0.11 | 12.8 ± 1.8 | 14.0 ± 2.6 |
| | Bound d < 0.5 nm | 0.91 ± 0.11 | 12.8 ± 1.8 | 14.0 ± 2.6 |
| | Bound c > 45 | 0.96 ± 0.12 | 12.8 ± 1.8 | 13.4 ± 2.5 |
| | Unbound d > 1.0 nm | 0.91 ± 0.11 | 13.7 ± 1.9 | 15.0 ± 2.8 |
| | Unbound d > 1.1 nm | 0.91 ± 0.11 | 13.1 ± 1.8 | 14.3 ± 2.7 |
| | Unbound c < 12 | 0.91 ± 0.11 | 15.3 ± 2.2 | 16.8 ± 3.2 |
| | Unbound c < 15 | 0.91 ± 0.11 | 17.2 ± 2.3 | 18.9 ± 3.4 |

*Table S12n. Kinetic parameters of EQVSAV A53G with the original and alternative COMMITTOR thresholds (see Table S1a for details).*

| Dataset | Thresholds | r_kon (p-value) | r_koff (p-value) | r_KD (p-value) |
|---|---|---|---|---|
| **EQVTAV** | Original | 0.14 (0.458) | 0.77 (0.064) | 0.97 (0.036) |
| | Bound, checked every 1 ps | 0.09 (0.485) | 0.77 (0.063) | 0.96 (0.041) |
| | Bound d < 0.45 nm | 0.14 (0.458) | 0.77 (0.063) | 0.97 (0.037) |
| | Bound d < 0.5 nm | 0.15 (0.455) | 0.77 (0.063) | 0.97 (0.037) |
| | Bound c > 45 | 0.12 (0.472) | 0.77 (0.064) | 0.96 (0.039) |
| | Unbound d > 1.0 nm | 0.14 (0.458) | 0.79 (0.050) | 0.96 (0.032) |
| | Unbound d > 1.1 nm | 0.14 (0.458) | 0.76 (0.065) | 0.96 (0.035) |
| | Unbound c < 12 | 0.14 (0.458) | 0.78 (0.058) | 0.97 (0.036) |
| | Unbound c < 15 | 0.14 (0.458) | 0.75 (0.077) | 0.96 (0.041) |

*Table S2a**.** Pearson correlation coefficient (r) between experimental and multi-eGO kon, koff and $K_D$ for each set of thresholds of Table S1, with p-value in parentheses, computed as described in the Methods. Correlations for $K_D$ were computed on ln K_D, as in Figure 5; small differences from the values in Figure 5 arise from the bootstrap resampling of the kinetic parameters. Changing the COMMITTOR threshold does not change the p-value significance of EQVTAV.*

| Dataset | Thresholds | r_kon (p-value) | r_koff (p-value) | r_KD (p-value) |
|---|---|---|---|---|
| **EQVSAV** | Original | 0.42 (0.295) | 0.30 (0.272) | 0.65 (0.120) |
| | Bound, checked every 1 ps | 0.47 (0.264) | 0.30 (0.275) | 0.62 (0.129) |
| | Bound d < 0.45 nm | 0.38 (0.316) | 0.30 (0.272) | 0.64 (0.120) |
| | Bound d < 0.5 nm | 0.39 (0.307) | 0.30 (0.271) | 0.62 (0.129) |
| | Bound c > 45 | 0.46 (0.269) | 0.30 (0.275) | 0.63 (0.122) |
| | Unbound d > 1.0 nm | 0.42 (0.294) | 0.21 (0.299) | 0.57 (0.151) |
| | Unbound d > 1.1 nm | 0.42 (0.293) | 0.25 (0.297) | 0.60 (0.139) |
| | Unbound c < 12 | 0.42 (0.295) | 0.19 (0.301) | 0.56 (0.156) |
| | Unbound c < 15 | 0.42 (0.293) | 0.12 (0.322) | 0.50 (0.183) |

*Table S2b. Correlation between experimental and multi-eGO kinetic parameters for EQVSAV with the original and alternative COMMITTOR thresholds (see Table S2a for details).*

| Dataset | Thresholds | r_kon (p-value) | r_koff (p-value) | r_KD (p-value) |
|---|---|---|---|---|
| **Pooled** | Original | 0.13 (0.405) | 0.47 (0.093) | 0.61 (0.040) |
| | Bound, checked every 1 ps | 0.13 (0.404) | 0.47 (0.093) | 0.59 (0.047) |
| | Bound d < 0.45 nm | 0.12 (0.418) | 0.47 (0.093) | 0.61 (0.041) |
| | Bound d < 0.5 nm | 0.14 (0.402) | 0.47 (0.093) | 0.61 (0.042) |
| | Bound c > 45 | 0.19 (0.363) | 0.47 (0.093) | 0.62 (0.040) |
| | Unbound d > 1.0 nm | 0.13 (0.404) | 0.42 (0.110) | 0.59 (0.044) |
| | Unbound d > 1.1 nm | 0.13 (0.405) | 0.45 (0.102) | 0.60 (0.042) |
| | Unbound c < 12 | 0.13 (0.402) | 0.39 (0.124) | 0.56 (0.054) |
| | Unbound c < 15 | 0.13 (0.406) | 0.32 (0.157) | 0.54 (0.065) |

*Table S2c. Correlation between experimental and multi-eGO kinetic parameters for the pooled data with the original and alternative COMMITTOR thresholds (see Table S2a for details).*

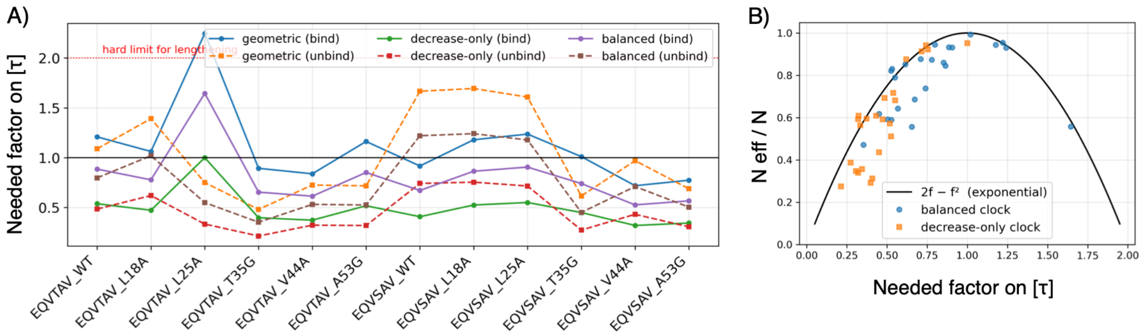


*Figure S1. Choice of the clock factor for the maximum-caliber reweighting. (A) Factor by which the simulated mean first-passage time of each system must change to reproduce experiment, f = A/c, for binding (solid lines) and unbinding (dashed lines) and three choices of the clock factor c: the geometric mean of the acceleration factors (geometric); the value for which all simulated times need to be shortened, f ≤ 1 (decrease-only); and the value that maximizes the smallest effective sample size (balanced), used in this work. Values f > 1 require lengthening the simulated times; for an exponential distribution this is only possible up to f = 2 (red dotted line), where the effective sample size vanishes. (B) Normalized effective sample size, N_eff/N, as a function of f for the balanced and decrease-only clocks. The solid line is the analytical result for reweighting an exponential distribution, N_eff/N = 2f − f².*

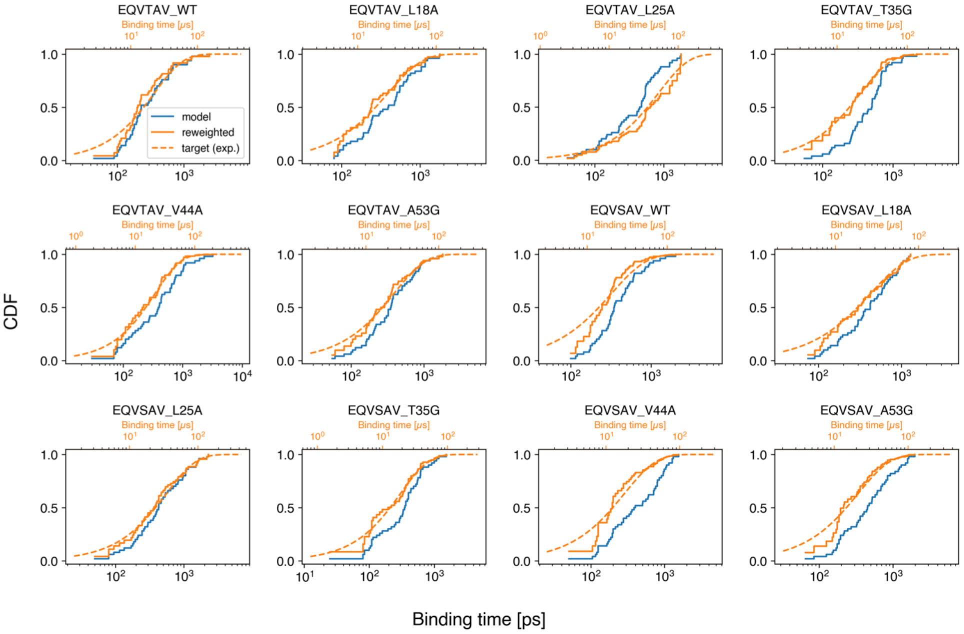


*Figure S2. Cumulative distributions of the binding times for each system. The distribution of the 50 simulated first-passage times (model, blue) is compared with the same distribution after maximum-caliber reweighting (reweighted, orange) and with the target exponential distribution (dashed), whose mean is the experimental binding time converted to simulation time with the clock factor c. The bottom axis reports the simulation time; the top axis reports the corresponding experimental time scale (t/c).*

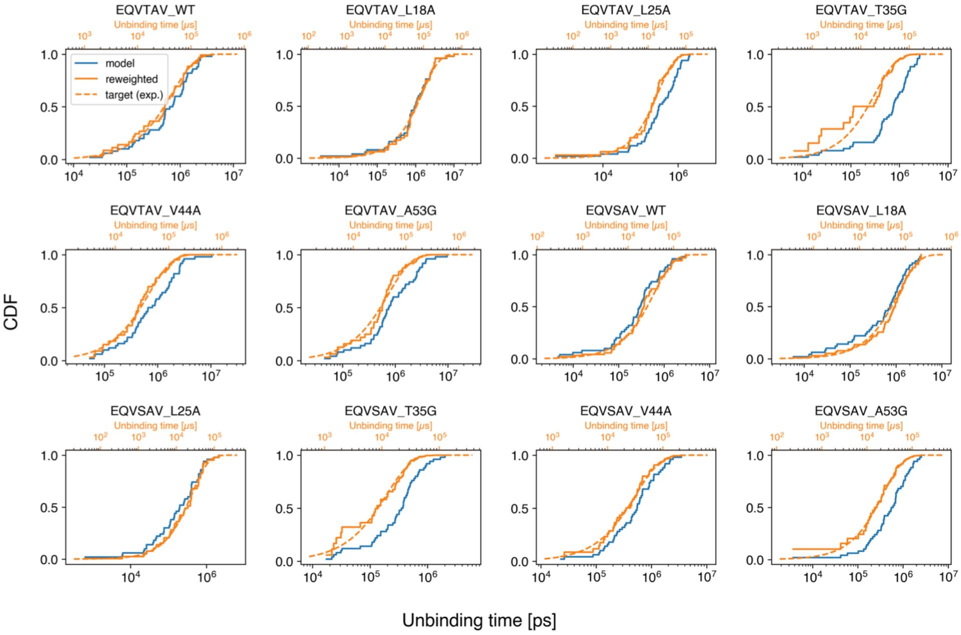


*Figure S3. Cumulative distributions of the unbinding times for each system. The distribution of the 50 simulated first-passage times (model, blue) is compared with the same distribution after maximum-caliber reweighting (reweighted, orange) and with the target exponential distribution (dashed), whose mean is the experimental unbinding time converted to simulation time with the clock factor c. The bottom axis reports the simulation time; the top axis reports the corresponding experimental time scale (t/c).*

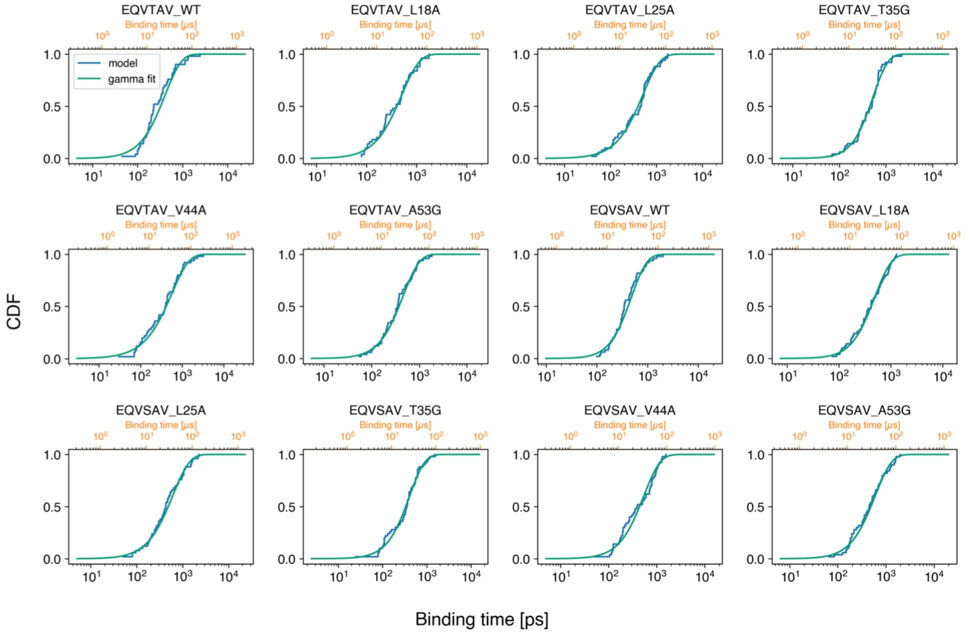


*Figure S4. Gamma fits of the simulated binding-time distributions used to estimate f_sim in the maximum-caliber weights. For each system, the cumulative distribution of the 50 simulated first-passage times (model, blue) is compared with the cumulative distribution of the fitted gamma distribution (green). The bottom axis reports the simulation time; the top axis reports the corresponding experimental time scale.*

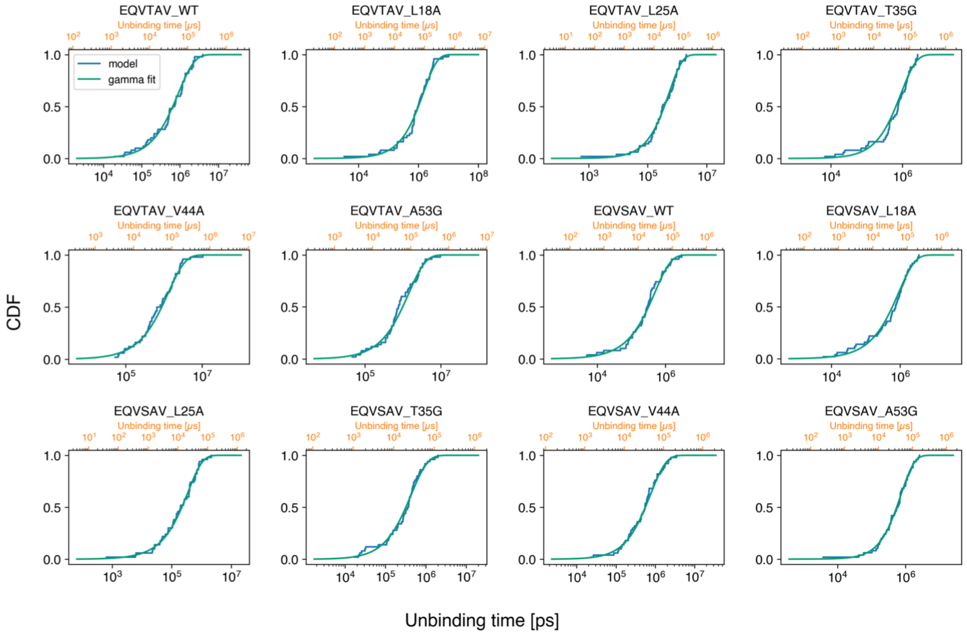


*Figure S5. Gamma fits of the simulated unbinding-time distributions used to estimate f_sim in the maximum-caliber weights. For each system, the cumulative distribution of the 50 simulated first-passage times (model, blue) is compared with the cumulative distribution of the fitted gamma distribution (green). The bottom axis reports the simulation time; the top axis reports the corresponding experimental time scale.*

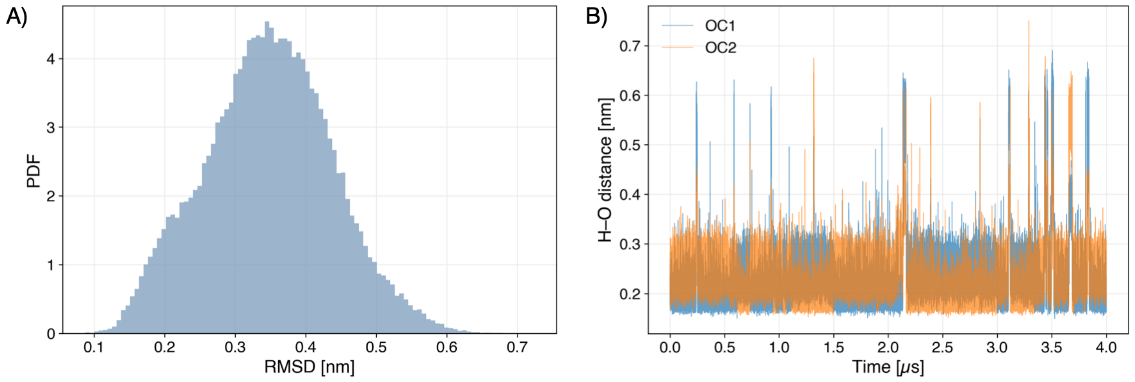


*Figure S6. Stability of the bound EQVTAV peptide in the reference simulations (four replicas, concatenated).* ***(A)*** *Distribution of the backbone RMSD of the bound peptide with respect to its initial pose, after fitting on PDZ2.* ***(B)*** *Time series of the distance between the backbone amide hydrogen of L25 and the two C-terminal carboxylate oxygens (OC1, OC2) of the bound peptide. The hydrogen bond is maintained throughout the simulations, alternately between the two oxygens.*

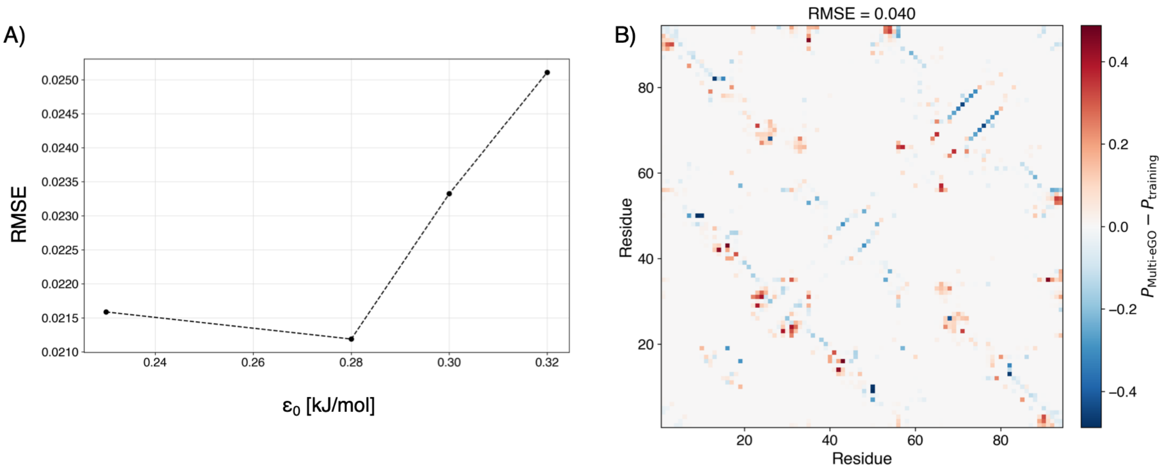


*Figure S7. Optimization of the intramolecular energy scale of the PDZ2 system. (A) Root-mean-square error (RMSE) between the reference and multi-eGO root mean square fluctuations (RMSF), excluding the first and last two residues, as a function of $\varepsilon_0^{intra}$. (B) Difference between the intramolecular contact probability maps of PDZ2 obtained from the multi-eGO ($\varepsilon_0^{intra}$ = 0.28 kJ/mol) and reference simulations. The RMSE over all residue pairs is reported above the map.*

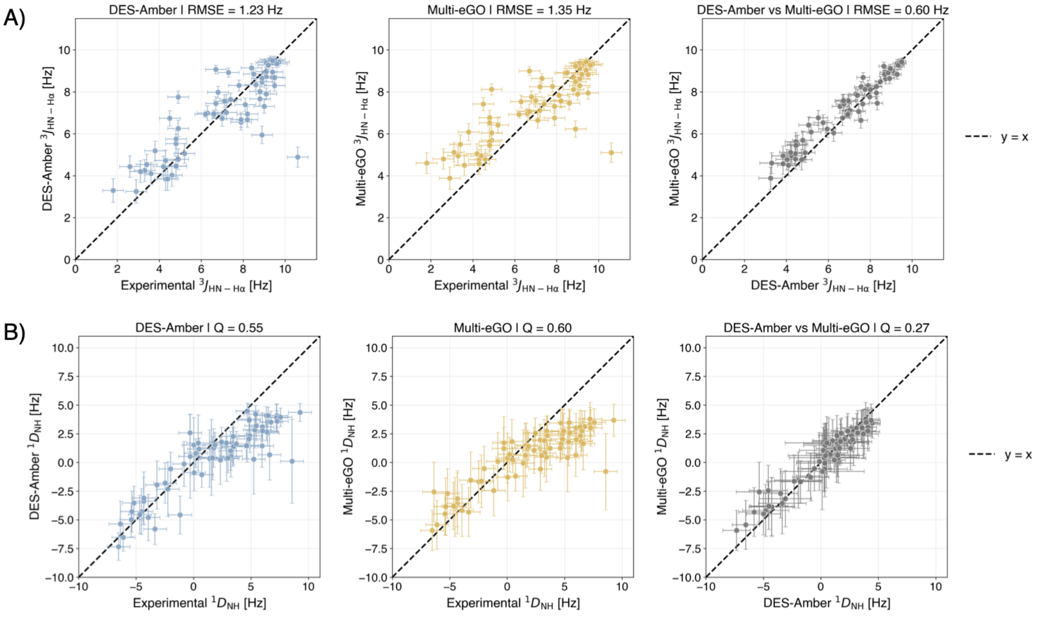


*Figure S8. Comparison with experimental NMR data for PDZ2 (BMRB 5131). (A) $^3J$(HN–Hα) scalar couplings computed from the DES-Amber (left) and multi-eGO (centre) simulations versus the experimental values, and comparison between the two models (right). Couplings were computed from the backbone φ angles of each frame with three Karplus parametrizations and then averaged over frames; simulation error bars are the standard deviation over the three parametrizations, and experimental error bars correspond to 0.5 Hz. (B) $^1D$(NH) residual dipolar couplings, computed with PALES (best-fit alignment tensor for each frame) and averaged over frames, compared in the same way. Agreement is quantified by the RMSE or the Q factor and reported above each panels, Q = [Σᵢ (D_calc,i − D_ref,i)² / Σᵢ D_ref,i²]^½, where the reference is the experimental data (left, centre) or DES-Amber (right) In the right panel, RMSE and Q are calculated using DES-Amber as the reference model to assess how accurately multi-eGO reproduces the DES-Amber ensemble. The low RMSE and Q values indicate good agreement between the two models.*

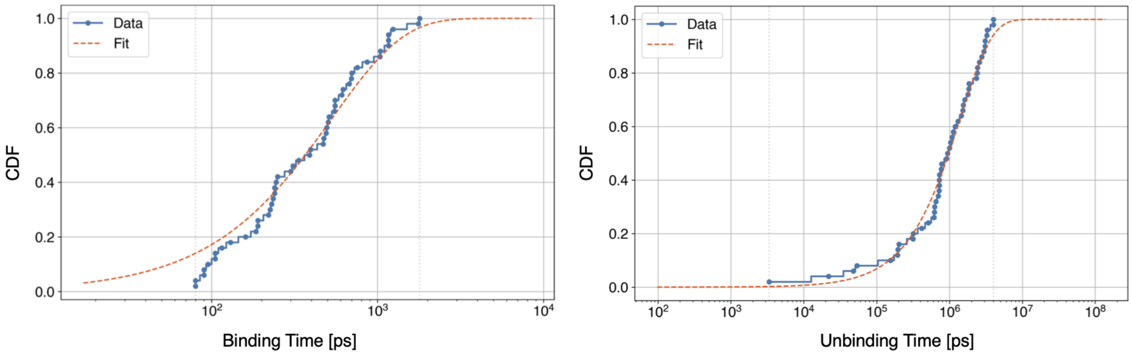


*Figure S9. Cumulative distribution function (CDF) of the PDZ2-L18A:EQVTAV binding/unbinding times (ps) obtained from 50 independent multi-eGO simulations, together with the corresponding exponential fit. Binding: p-value = 0.25, tau = 524 ± 89 ps. Unbinding p-value = 0.53, tau = 1,401,000 ± 257,000 ps.*

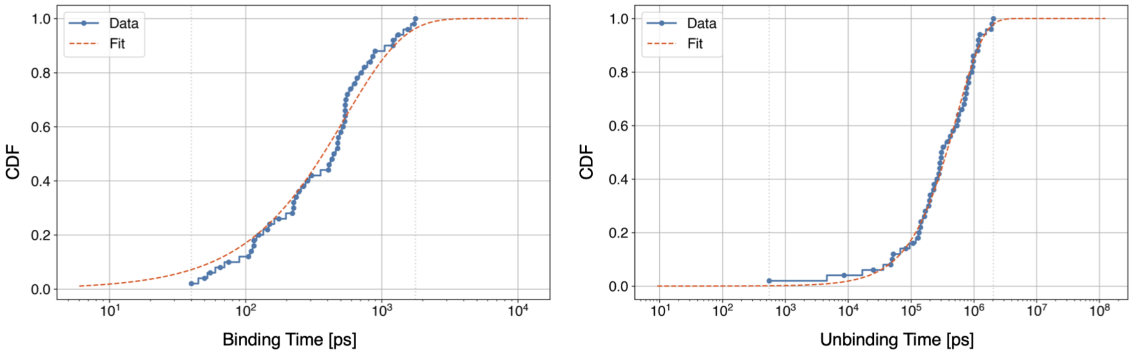


*Figure S10. Cumulative distribution function (CDF) of the PDZ2-L25A:EQVTAV binding/unbinding times (ps) obtained from 50 independent multi-eGO simulations, together with the corresponding exponential fit. Binding: p-value = 0.55, tau = 533 ± 93 ps. Unbinding p-value = 0.94, tau = 528,000 ± 114,000 ps.*

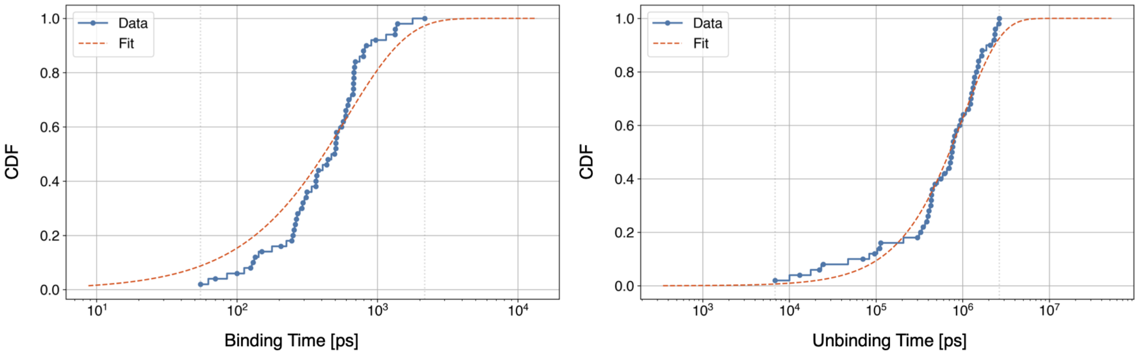


*Figure S11. Cumulative distribution function (CDF) of the PDZ2-T35G:EQVTAV binding/unbinding times (ps) obtained from 50 independent multi-eGO simulations, together with the corresponding exponential fit. Binding: p-value = 0.08, tau = 592 ± 80 ps. Unbinding p-value = 0.71, tau = 986,000 ± 161,000 ps.*

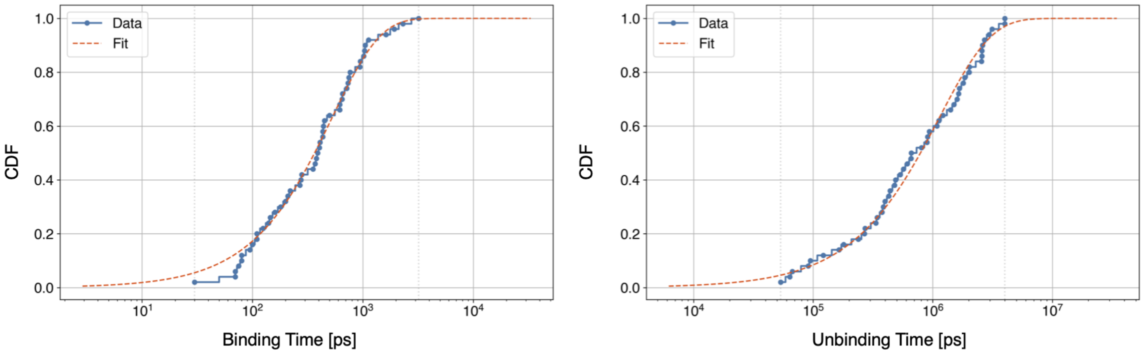


*Figure S12. Cumulative distribution function (CDF) of the PDZ2-V44A:EQVTAV binding/unbinding times (ps) obtained from 50 independent multi-eGO simulations, together with the corresponding exponential fit. Binding: p-value = 0.61, tau = 523 ±*

*111 ps. Unbinding p-value = 0.93, tau = 1,123,000 ± 260,000 ps.*

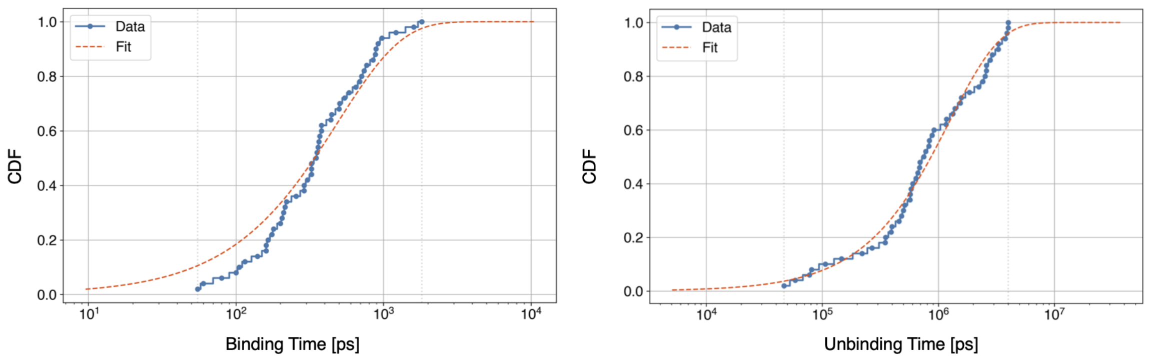


*Figure S13. Cumulative distribution function (CDF) of the PDZ2-A53G:EQVTAV binding/unbinding times (ps) obtained from 50 independent multi-eGO simulations, together with the corresponding exponential fit. Binding: p-value = 0.27, tau = 493 ± 74 ps. Unbinding p-value = 0.73, tau = 1,254,000 ± 289,000 ps.*

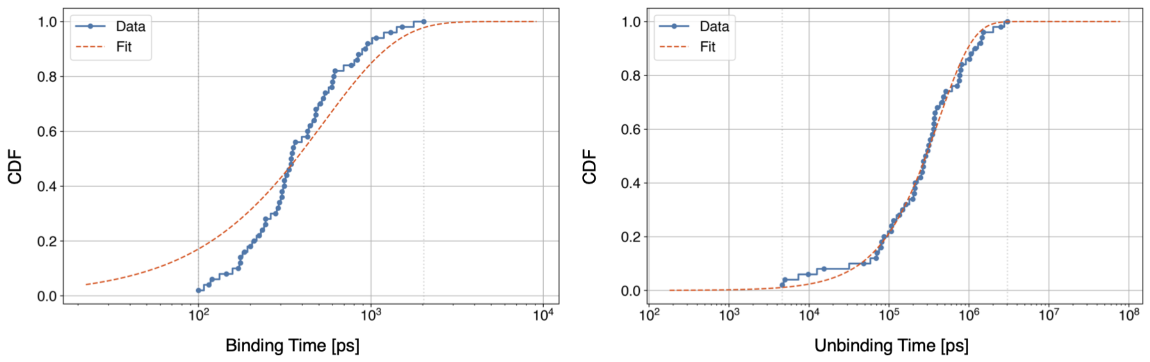


*Figure S14. Cumulative distribution function (CDF) of the PDZ2-WT:EQVSAV binding/unbinding times (ps) obtained from 50 independent multi-eGO simulations, together with the corresponding exponential fit. Binding: p-value = 0.04, tau = 525 ± 64 ps. Unbinding p-value = 0.90, tau = 415,000 ± 95,000 ps.*

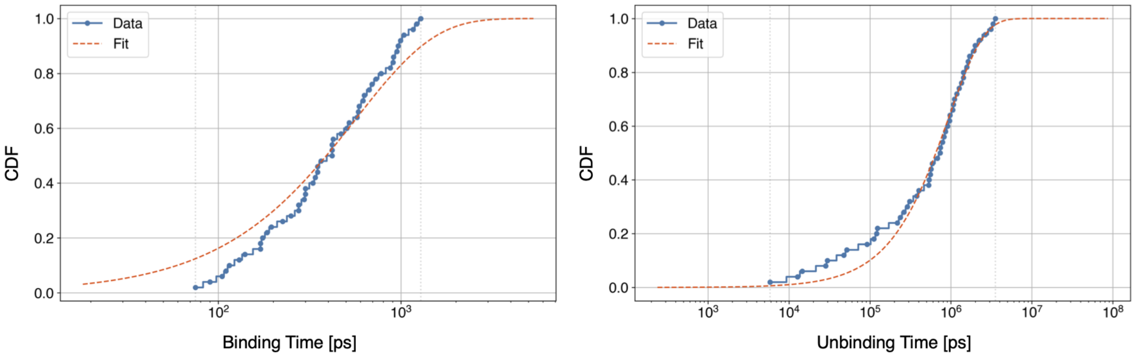


*Figure S15. Cumulative distribution function (CDF) of the PDZ2-L18A:EQVSAV binding/unbinding times (ps) obtained from 50 independent multi-eGO simulations, together with the corresponding exponential fit. Binding: p-value = 0.34, tau = 560 ± 80 ps. Unbinding p-value = 0.70, tau = 924,000 ± 199,000 ps.*

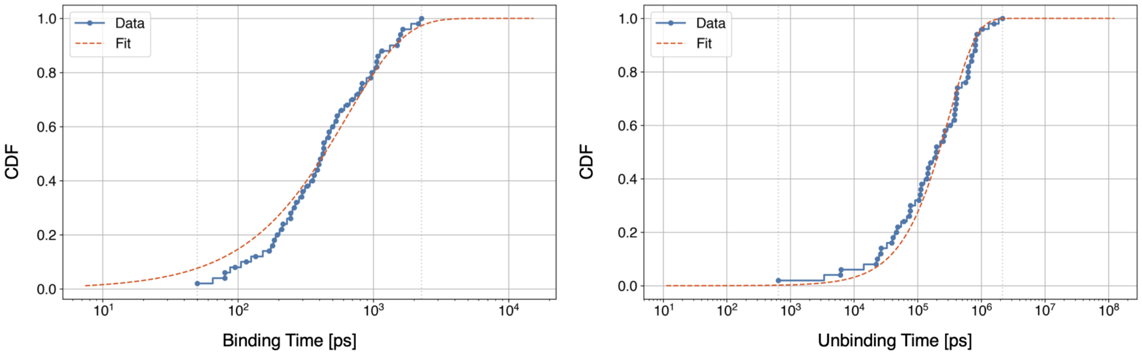


*Figure S16. Cumulative distribution function (CDF) of the PDZ2-L25A:EQVSAV binding/unbinding times (ps) obtained from 50 independent multi-eGO simulations, together with the corresponding exponential fit. Binding: p-value = 0.47, tau = 612 ± 105 ps. Unbinding p-value = 0.63, tau = 309,000 ± 86,000 ps.*

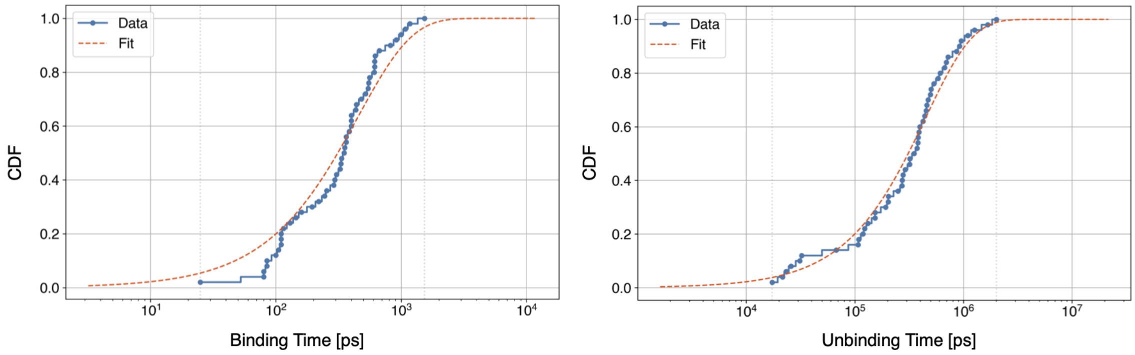


*Figure S17. Cumulative distribution function (CDF) of the PDZ2-T35G:EQVSAV binding/unbinding times (ps) obtained from 50 independent multi-eGO simulations, together with the corresponding exponential fit. Binding: p-value = 0.23, tau = 446 ± 69 ps. Unbinding p-value = 0.72, tau = 437,000 ± 73,000 ps.*

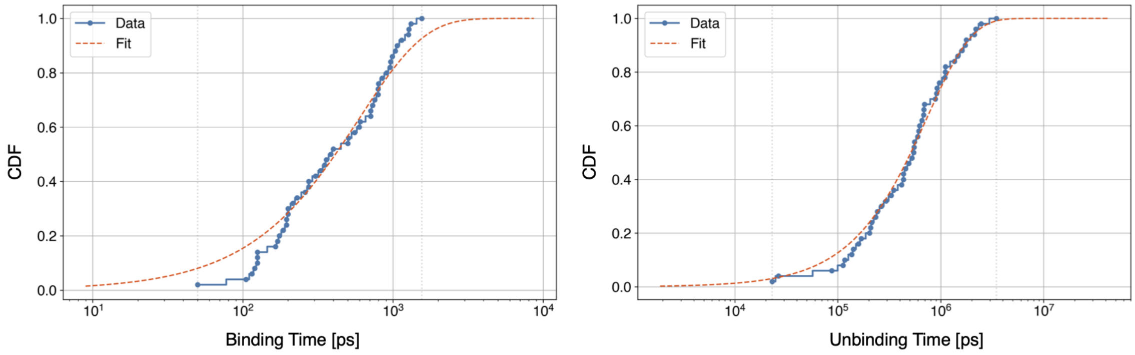


*Figure S18. Cumulative distribution function (CDF) of the PDZ2-V44A:EQVSAV binding/unbinding times (ps) obtained from 50 independent multi-eGO simulations, together with the corresponding exponential fit. Binding: p-value = 0.25, tau = 595 ± 94 ps. Unbinding p-value = 0.87, tau = 738,000 ± 143,000 ps.*

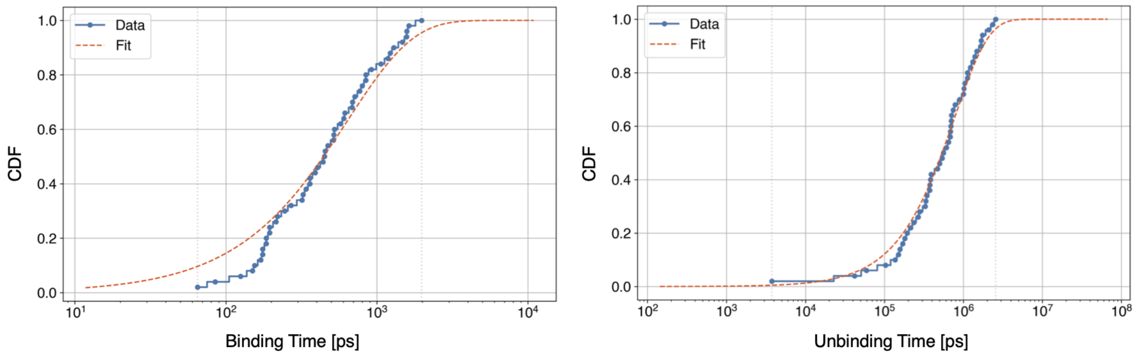


*Figure S19. Cumulative distribution function (CDF) of the PDZ2-A53G:EQVSAV binding/unbinding times (ps) obtained from 50 independent multi-eGO simulations, together with the corresponding exponential fit. Binding: p-value = 0.20, tau = 637 ± 106 ps. Unbinding p-value = 0.87, tau = 765,000 ± 140,000 ps.*

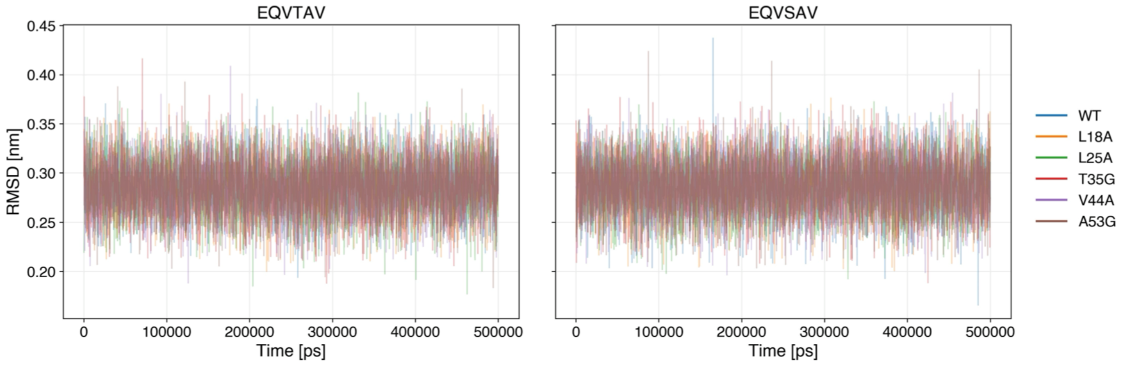


*Figure S20. RMSD of PDZ2 as a function of time for each multi-eGO system simulation considering the different protein mutations, with the EQVTAV (left) and EQVSAV (right) peptides, computed with respect to the starting structure (PDZ2 with the bound peptide). In all systems the RMSD remains stable at about 0.25–0.30 nm, indicating that the WT and all mutants remain folded in the bound state.*

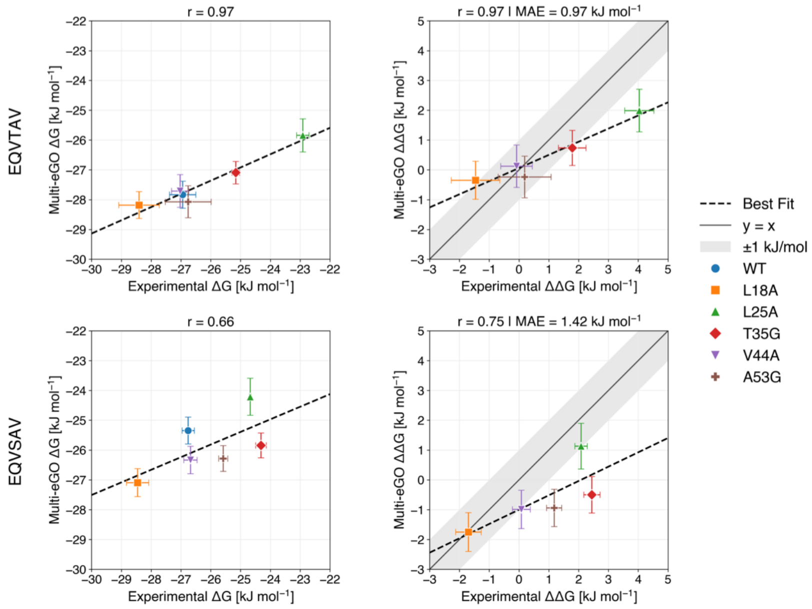


*Figure S21. Binding free energies (left) and changes upon mutation (right) for the EQVTAV (top) and EQVSAV (bottom) peptides. $\Delta G = RT \ln K_D$ was computed from the experimental (277 K) and multi-eGO (283 K) dissociation constants, and $\Delta\Delta G = \Delta G(\text{mutant}) - \Delta G(\text{WT})$ with respect to the WT complex of the same peptide; errors are propagated from those on $K_D$. Dashed lines are least-squares linear fits; in the ΔΔG panels, the solid line indicates y = x and the shaded area deviations within ±1 kJ/mol. The Pearson correlation coefficient (r) and, for ΔΔG, the mean absolute error (MAE) are reported above each panel.*
